# A Data-Driven Image Extraction and Analysis Pipeline for Plant Phenotyping in Controlled Environments

**DOI:** 10.64898/2026.02.25.707797

**Authors:** Fahimeh Orvati Nia, Joshua Peeples, Seth C. Murray, Andrew McFarland, Troy Vann, Shima Salehi, Robert Hardin, David D. Baltensperger, Amir M.H. Ibrahim, J. Alex Thomasson, Henry Fadamiro, Nithya K. Subramanian, Suresh D. Pillai, Rebecca Roston, Joslin Ishimwe, Diptadeep Basak, Nazar Oladepo, Uday Vysyaraju

## Abstract

Advances in automation, imaging, and artificial intelligence have enabled large-scale plant phenotyping, but image analysis remains a critical bottleneck for crop improvement and biological discovery. We developed an integrated multispectral phenotyping framework using imagery from the Texas A&M AgriLife Precision Automated Phenotyping Greenhouse and expanded Plant Growth and Phenotyping (PGP v2) data across maize, cotton, rice, and sorghum. The pipeline integrates pseudo-RGB generation, plant detection and segmentation, image stitching, vegetation-index analysis, texture analysis, morphological trait extraction, and temporal comparison of image-derived features to quantify changes in plant structure, spectral reflectance, and texture over time. Among the evaluated segmentation approaches, SAM v3 provided the highest and most consistent accuracy across diverse crop structures, although it required greater computational time than classical methods. SAM2Long maintained plant-instance associations across vertically stacked frames, while Scale-Invariant Feature Transform (SIFT)-based stitching reconstructed plant mosaics when individual plants extended beyond a single field of view. For each plant and imaging date, the pipeline generated an 863-dimensional feature vector spanning vegetation indices, spectral statistics, texture descriptors, and morphological traits. The framework was evaluated through two case studies: treatment-level temporal analysis of mutagenized sorghum lines and cold-stress phenotyping of maize using a separate imaging system. In both studies, the extracted features supported statistical and multivariate analyses of phenotypic variation and enabled separation of plants based on treatment- or stress-related responses. The combined dataset and workflow provide structured, automated, and well-documented phenotypic analysis across multiple crops, experimental settings, and imaging systems for controlledenvironment plant science and crop improvement.

**Plain Language Summary:** Temporal imaging of plants in controlled environments helps scientists better understand growth and biological processes. However, analyzing large volumes of images has been limited by a lack of automated tools. Multispectral imagery captures additional information about plant pigments, structure, and stress beyond standard color images. We developed an automated analysis pipeline that identifies individual plants, tracks their growth over time, and measures traits such as height, area, shape, texture, and vegetation indices. Using artificial intelligence, the system efficiently processes thousands of images to provide consistent and repeatable measurements. By integrating engineering and plant biology, this work supports data-driven decisions for crop improvement and agricultural research.

## 1 Introduction

Significant investments in high-throughput plant phenotyping (HTP) controlled environment infrastructure have been made worldwide, leading to the rapid expansion of platforms such as automated greenhouses, growth chambers, and climate rooms across North America, Europe, Asia, and Australia (Rosenqvist et al., 2019). International surveys identify an extensive network of phenotyping facilities currently in operation globally, with a major focus on controlled environment systems designed for automated imaging under precisely regulated conditions (Yang et al., 2020). Although these facilities reflect a substantial global commitment to data-driven crop improvement, their full scientific potential has not been achieved, given recent advances in genomics, sensor engineering, and automated data collection (Fiorani and Schurr, 2013; Pieruschka and Schurr, 2019; Poorter et al., 2023). A recurring challenge has been the lack of coordinated data analytics, metadata management, and computational consistency to match advances in imaging hardware. Many early phenotyping programs relied heavily on project-specific analytical workflows, which limited reusability, made cross-study comparison difficult, and constrained long-term scalability (Schnaufer and Pistorius, 2020). Metadata structures were often inconsistent across institutions, and traits extracted from similar experimental designs could not be reliably compared due to the absence of shared descriptors and harmonized documentation (Papoutsoglou et al., 2020).

To address these limitations, the phenotyping community has introduced a series of standardization and interoperability frameworks. Among them, the Minimum Information About a Plant Phenotyping Experiment (MIAPPE) initiative provides structured guidelines for experiment description, controlled vocabularies, and metadata schemas aligned with the Findable, Accessible, Interoperable, and Reusable (FAIR) principles (Reiser et al., 2018; Papoutsoglou et al., 2020), which informed the design goals of the present framework. These developments underscore a broader consensus that advances in controlled environment phenomics require more than imaging innovation alone. Progress also depends on robust software infrastructure, transparent data workflows, well-documented computation, and sustained interaction across engineering, computational, and plant science disciplines (Poorter et al., 2023; Pieruschka and Schurr, 2019; Tripodi et al., 2022). Contemporary analyses emphasize that biological discovery is facilitated when imaging pipelines, experimental metadata, and analytical modules are co-developed and maintained through coordinated program management rather than through isolated technical efforts (Fiorani and Schurr, 2013; Poorter et al., 2023). As controlled environment platforms continue to scale in complexity and throughput, these principles form the foundation for creating systems that can produce reliable, interpretable, and comparable phenotypic information across large spatial and temporal scales.

Within this context, the Texas A&M AgriLife Automated Precision Phenotyping Greenhouse provides an example of a controlled environment system structured around continuous coordination between technical and biological teams. The facility integrates version-controlled repositories, shared preprocessing pipelines, and standardized metadata management within the implementation, enabling iterative refinement of analytical workflows while maintaining consistency across experiments. Building on this organizational foundation, this work introduces an end-to-end automated controlled environment phenotyping framework unifying multispectral image acquisition, calibration, segmentation, instance tracking, and trait extraction within a modular and well-documented software architecture. Leveraging the Plant Growth and Phenotyping (PGP) v1 dataset (Zambre et al., 2024), an updated PGP v2 dataset, reported here, further expands the system to additional crops, treatments, and spectral modalities for algorithmic development. The entire PGP v2 dataset across crops was used to build the analytical framework presented throughout. The analytical framework is then demonstrated through two example case studies: one on sorghum (*Sorghum bicolor* L.) using PGP v2 data collected in the automated greenhouse, and a second using an independently collected maize (*Zea mays L.) dataset acquired with a standard RGB camera. Together, the two case studies highlight the utility of the framework for treatment-level temporal analysis and stress-response phenotyping across different crop experiments and imaging systems.* Through this design, the framework demonstrates that coordinated management, harmonized analytics, and controlled environment imaging can collectively support scalable, reliable, and biologically meaningful phenotyping in modern controlled environment programs. The analytical framework is open-source and available to other researchers, both through a web-based interface for direct plant-scientist use and as code for future tool development.

## 2 Materials and Methods

### 2.1 Facility Overview

The Texas A&M AgriLife Automated Precision Phenotyping Greenhouse (APPG) (Thomasson et al., 2022) is a controlled environment research facility developed to support high-throughput plant phenotyping under reproducible environmental conditions. The greenhouse infrastructure is comprised of modular growth zones equipped with automated systems for regulating temperature, humidity, light intensity, and photoperiod. Environmental parameters are centrally managed through an integrated control system that enables precise scheduling and spatial uniformity across treatment groups. A robotic gantry mechanism allows for repeatable plant access along X, Y, and Z axes, facilitating noninvasive imaging and sensor deployment. Plants are arranged on stationary benching systems aligned for efficient gantry traversal. Supporting infrastructure includes a headhouse for pot preparation and sensor maintenance, as well as adjacent workstations for experiment monitoring and data review. These features enable long-term, multi-crop experiments with scalable data collection and robust environmental consistency. The facility consists of five interconnected greenhouse bays. Individual greenhouse bays range from approximately 558 to 586 m^2^ in area, including an acclimatization yard (558 m^2^) and four primary production bays (558 m^2^, 560 m^2^, 558 m^2^, and 586 m^2^). In total, the building comprises 3,970 m^2^ of gross area (3,793 m^2^ net usable area), supporting large-scale and multi-treatment crop experiments under controlled conditions (McFarland et al., 2025).

### 2.2 Project Organization and Workflow Integration

Figure 1 outlines the data workflow associated with the APPG. The analytical project team was organized into four functional groups: data collection, data preprocessing, data analysis, and data discovery. The current core team included eight members: two responsible for data collection, four contributing to data preprocessing and analysis, and two supporting data discovery. Additional plant science collaborators contributed to individual case studies. Plant science collaborators contributed experiments and longitudinal imaging datasets that created practical analytical needs for pipeline development. In this bidirectional collaboration, plant science researchers provided biologically relevant imaging problems that could not be easily analyzed with existing tools, while the engineering and analytics teams developed processing workflows to return structured phenotypic outputs for those experiments. Greenhouse operations personnel manage plant care, environmental regulation, and infrastructure maintenance. Imaging specialists oversee data acquisition, sensor calibration, and imaging schedules. Data preprocessing teams perform image normalization, segmentation, and trait isolation. Analytics teams conduct statistical evaluation and visualization. Data discovery researchers investigate trait patterns, genotype-phenotype associations, and emergent biological signals. The analysis and discovery stages form an iterative feedback loop in which insights from data discovery inform subsequent analyses until the desired end product is reached. The Administration team facilitates project oversight, resource management, and interdisciplinary communication to ensure sustained and well-documented outcomes. An overarching goal of integrating these teams was to use each study to maximize the number and types of generalized phenotypic features extracted, applicable across species, and to output useful measures for future studies of interest. A long-term goal of the framework is to mature into a system that enables plant science researchers to process longitudinal imaging experiments independently and translate those data into biological knowledge with minimal technical friction.

**Figure 1:**
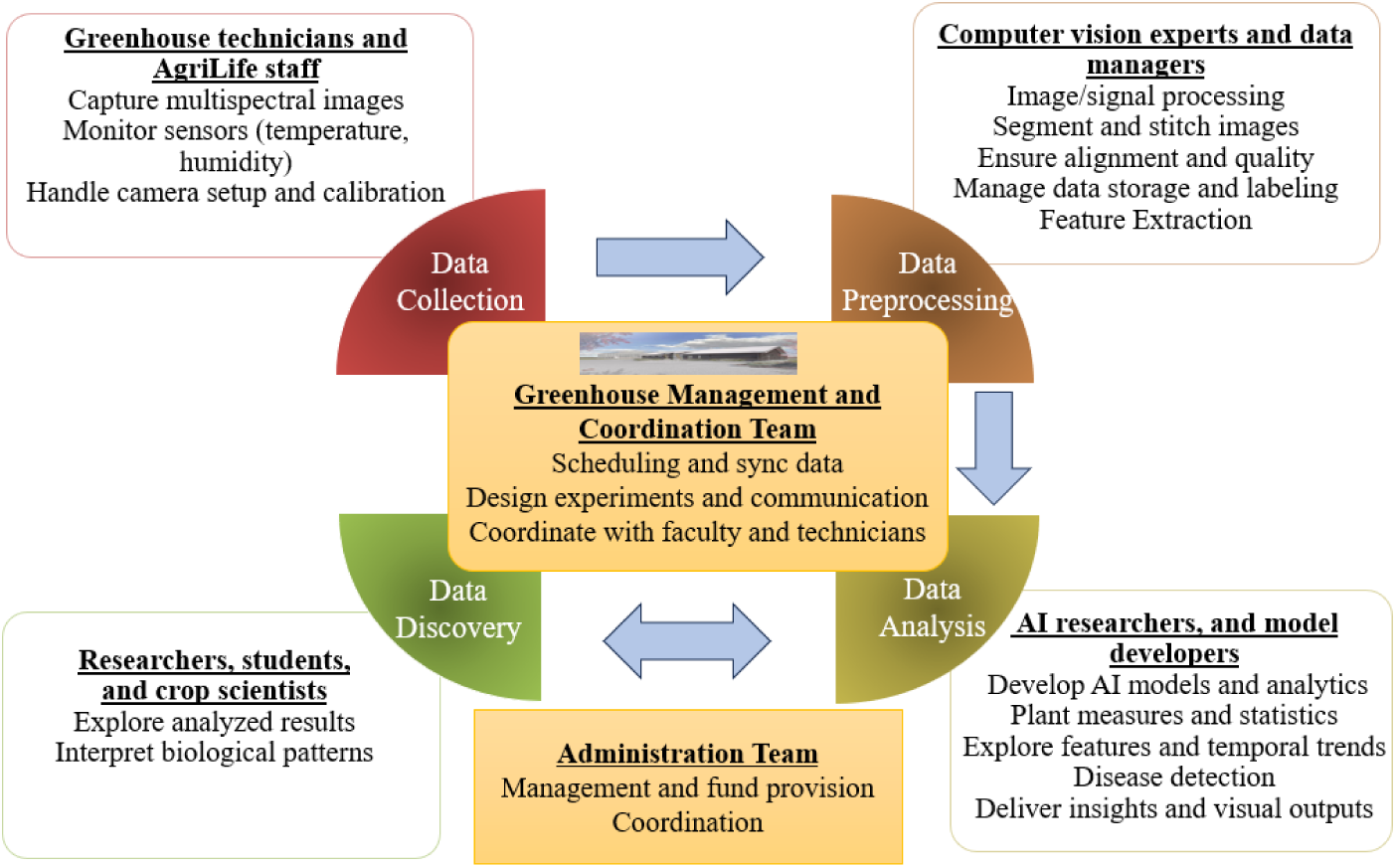
Integration of teams and responsibilities in the Texas A&M AgriLife Automated Precision Phenotyping Greenhouse (APPG). The bidirectional arrow between Data Analysis and Data Discovery represents the iterative feedback loop in which insights from data discovery inform subsequent analyses.

### 2.3 Multispectral Data Acquisition and Dataset Expansion

Plants were imaged on fixed benches arranged in parallel rows to form a regular grid across each greenhouse bay. During an imaging run, the robotic gantry traversed the greenhouse by moving longitudinally along bench aisles and stepping laterally between adjacent rows, enabling repeatable coverage of all plants in a row-by-row pattern (Figure 2a). At each imaging position, the camera head was positioned above the canopy at a consistent working distance and captured a vertically stacked sequence of frames, defined as consecutive images acquired at camera positions along the Z-axis from the base to the canopy apex of each plant. This acquisition strategy provided aligned multispectral snapshots across the full plant profile.

**Figure 2:**
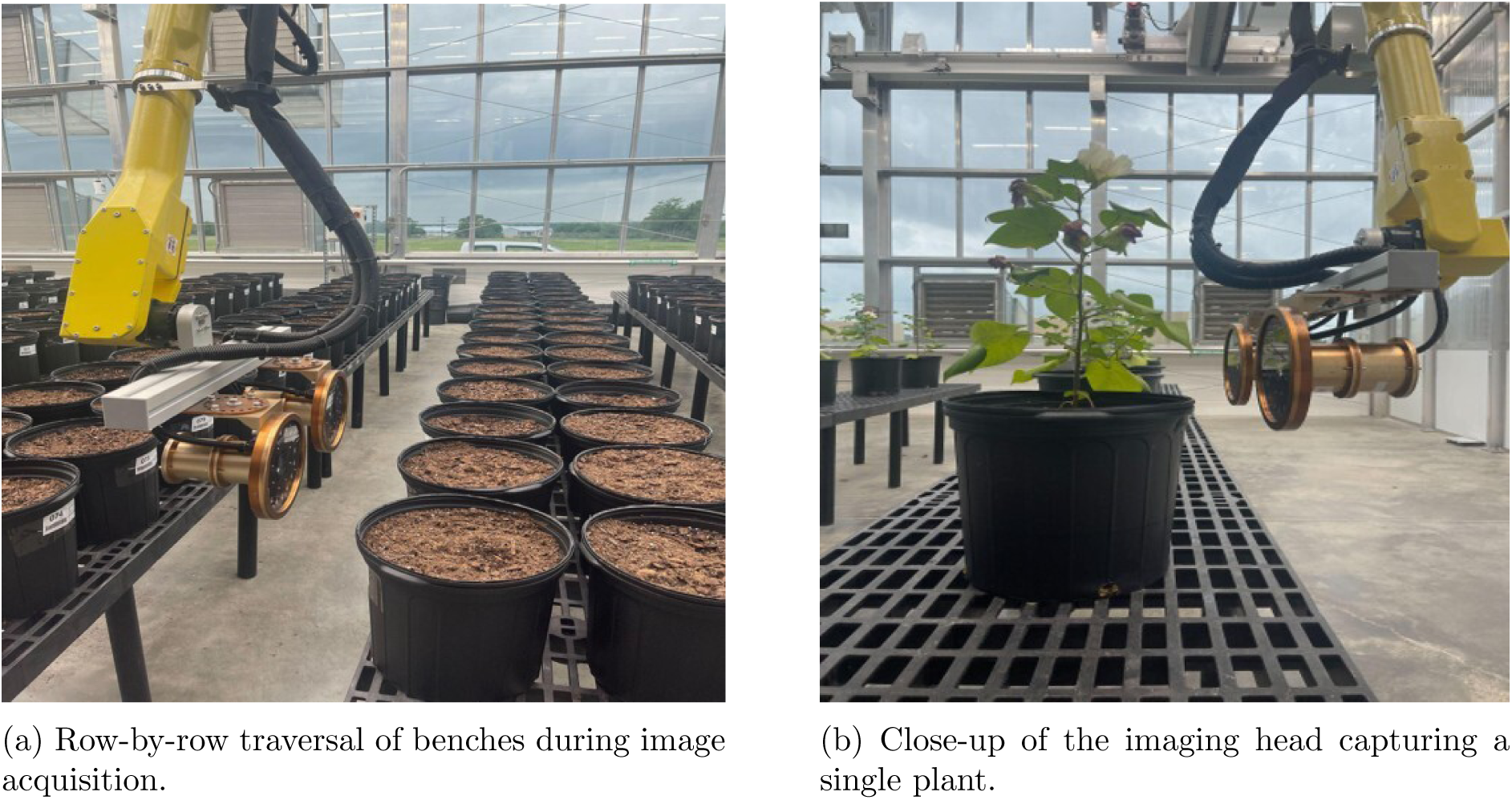
Robotic multispectral image acquisition in the Texas A&M AgriLife APPG.

In the Texas A&M AgriLife APPG data collection platform, the camera height is manually adjusted by the operator prior to each imaging session based on the approximate canopy height of the plants being imaged. During each imaging session, the robotic gantry maintains a consistent working distance between the camera and plant canopy for all frames captured within that session. Although this distance may vary between sessions due to manual adjustment, it remains effectively constant within each session, supporting stable imaging geometry for temporal analysis. Because the camera-to-plant distance may vary between imaging sessions, pixel-based morphological measurements may also vary in scale. Therefore, these measurements were converted to physical units using the recorded positional metadata and known camera parameters, as described in Section 2.5.2.

Together, the row-by-row gantry traversal, vertical frame acquisition, and recorded positional metadata enable longitudinal comparison of individual plants within and across APPG experiments (Figure 2b). For each image acquisition, the greenhouse system automatically records metadata including species, room number, date, time, image index, row number, plant number, orientation, and the spatial coordinates of both the plant and the robot. These metadata are stored in a structured log file and linked to each image, enabling automated spatial calibration, plant identification, and temporal tracking across imaging sessions without manual annotation. An example of the metadata structure is provided in Supplemental Table S1.

APPG multispectral imaging was performed using the MSISAGRI1A multispectral camera (Zambre et al., 2024), which integrates a 4-megapixel CMOS sensor with a synchronized four-channel LED illumination system. The four spectral bands (yellow, 580 nm; red, 660 nm; red-edge, 735 nm; and near-infrared, 820 nm) capture information related to chlorophyll concentration, canopy structure, and plant stress. The camera captures all bands simultaneously in snapshot mode using AntiXTalk^TM^ technology to minimize inter-band leakage and preserve spectral fidelity. Each band is stored as a separate channel within a 16-bit TIFF image of 512 × 512 pixels, providing consistent spatial resolution across bands. The system operates at up to 180 frames s*^−^*^1^ and is controlled via a water-resistant embedded PC interface. APPG imaging sessions were conducted periodically between 2023 and 2025 across multiple species, enabling the capture of temporal changes in plant morphology and reflectance across developmental stages. Building on this acquisition framework, the expanded PGP version 2 dataset is introduced, extending the initial release (Zambre et al., 2024). The original PGP v1 dataset contained 1,137 images across maize, cotton, and rice. PGP v2 substantially increases both scale and diversity, comprising approximately 14,448 images of maize (*Zea mays* L.), 27,660 of cotton (*Gossypium hirsutum*), 688 of rice (*Oryza sativa*), and 10,608 of sorghum (*Sorghum bicolor*), along with 1,840 manually annotated sorghum images for keypoint detection. A detailed summary of the dataset, including the number of unique plants, imaging sessions, and frames per plant, is provided in Table 1.

**Table 1:** Summary of the PGP v2 dataset.

| Crop | Date Range | Sessions | Frequency | # Plants | Frames/Plant | Total Images |
| --- | --- | --- | --- | --- | --- | --- |
| Maize | Dec 2023–Jul 2024 | 40 | 3–4×/week | 271 | 6–15 | 14,448 |
| Cotton | Apr 2024–Jul 2024 | 66 | daily | 12 | 5–10 | 27,660 |
| Sorghum | Dec 2024–May 2025 | 17 | weekly | 48 | 13 | 10,608 |
| Rice | May 2024–Aug 2024 | 8 | weekly | 8 | 5–13 | 688 |

The dataset integrates multispectral imagery collected over multiple sessions, with each plant represented by a vertical sequence of frames acquired at successive camera positions along the Zaxis from the plant base to the canopy apex. The number of frames required for each plant varies across species and growth stages because of differences in plant height. Taller plants therefore require more vertically stacked frames to capture the full plant profile. This acquisition strategy provides multiple overlapping views of each plant along the canopy axis for subsequent tracking, stitching, and trait extraction. Data are organized hierarchically by species, imaging date, and frame index to facilitate temporal analysis and comparison across species based on temporal and structural imaging characteristics. Detailed treatment and genotype annotations are not uniformly available across all components of the dataset. Where available, such information is recorded in the associated metadata; however, the primary focus of this dataset description is on the imaging structure, acquisition protocol, and temporal sampling characteristics. The images are also broadly valuable for deep-learning algorithm development.

### 2.4 Data Processing Pipeline

An overview of the complete image processing pipeline is provided in Figure 3, illustrating each step from raw data acquisition to derived phenotypic features. This example is from the APPG, but other image collection approaches would work similarly. The complete pipeline is publicly available (Data Availability), and the workflow is designed to facilitate consistent and structured processing through standardized configuration files and data organization. Input images are organized by acquisition date and plant identifier, enabling automated end-to-end processing from raw multispectral data to derived phenotypic outputs.

**Figure 3:**
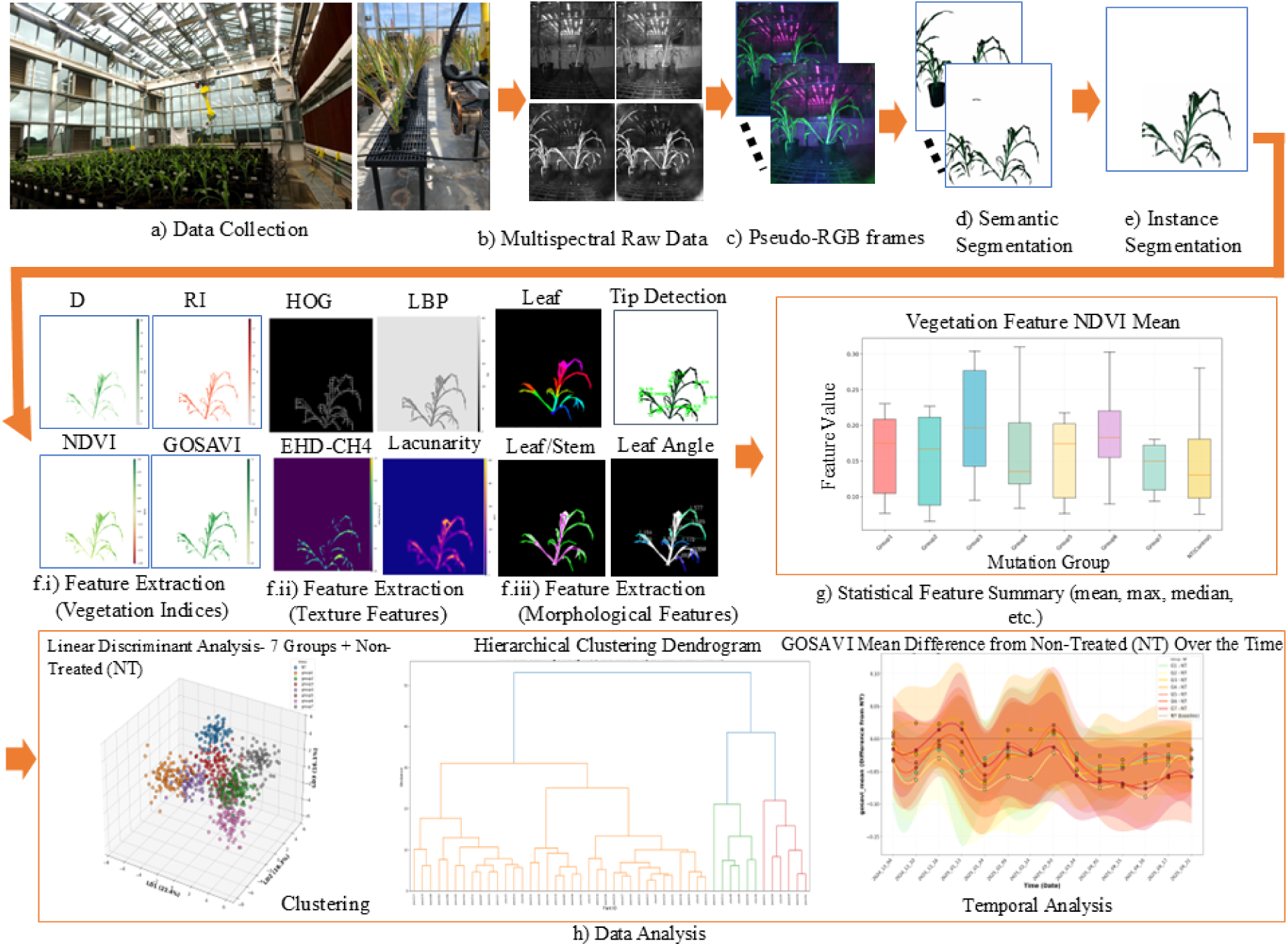
Comprehensive pipeline for multispectral plant phenotyping and feature analysis. (a) Data collection example inside the automated greenhouse (APPG). (b) Acquisition of four-band multispectral images (green, red, red-edge, near-infrared) from different views of each plant across multiple frames. (c) Generation of pseudo-RGB frames. (d) Semantic segmentation to remove background. (e) Instance segmentation to identify and separate individual plants. (f) Feature extraction examples including (i) vegetation indices (e.g., Difference Vegetation Index (DVI), Redness Index (RI), Normalized Difference Vegetation Index (NDVI), Green Optimized Soil-Adjusted Vegetation Index (GOSAVI)), (ii) texture descriptors (e.g., Histogram of Oriented Gradients (HOG), Local Binary Pattern (LBP), Edge Histogram Descriptor (EHD), Local Lacunarity), and (iii) morphological traits (e.g., leaf segmentation, tip detection, curvature angle). (g) Representative statistical feature summaries (e.g., mean, maximum, median) shown as box plots of NDVI mean across treatment groups. (h) Example data analysis outputs, including Linear Discriminant Analysis (LDA) (Zhao et al., 2024) for supervised multivariate visualization, hierarchical clustering for unsupervised grouping, and temporal analysis showing GOSAVI mean difference from the non-treated (NT) control over time.

#### 2.4.1 Image Preparation and Segmentation

To ensure compatibility with computer vision models trained on natural color imagery, multispectral bands were converted to 8-bit pseudo-RGB composites. Since the camera does not provide a blue band or a conventional green band, the 580 nm yellow/yellow-green channel was used as a surrogate visible band for pseudo-RGB generation and is referred to as *I*_green_ in the following equations for consistency with the implemented pipeline notation. Each spectral band image *I_λ_*, where *λ* ∈ {green, red, red-edge, near-infrared}, was normalized to the 0–255 intensity range according to Equation 1 (Patro and Sahu, 2015):

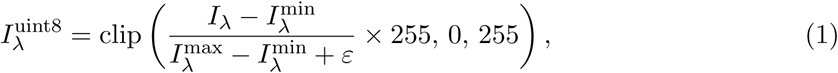

where *I_λ_*^min^ and *I_λ_*^max^ denote the minimum and maximum pixel intensities of the spectral band, and *ε* = 10*^−^*^6^ prevents division by zero. The clip operator ensures that intensities remain within the valid range and reduces the effect of noise or illumination variability. Per-band min– max normalization was applied independently to each spectral band to place all channels on a common 8-bit intensity scale for downstream analysis. The resulting arrays were cast into unsigned 8-bit format for compatibility with image processing and deep learning models. After normalization, the spectral bands were stacked to form a pseudo-RGB composite as shown in Equation 2 (Yang et al., 2020):

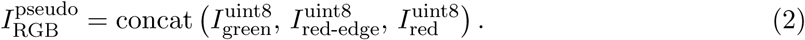

The near-infrared band could alternatively be used as a pseudo-RGB channel, for example by replacing the red-edge band, but was retained here for vegetation index computation. This preprocessing step rescaled the multispectral bands to a common 8-bit intensity range and enabled compatibility with convolutional neural networks pretrained on RGB imagery (Kattenborn et al., 2021). It did not constitute radiometric or color calibration across imaging sessions.

Plant segmentation was performed using a user-selectable model integrated within the phenotyping pipeline, with optional plant detection used to guide segmentation when required. Prior to adopting deep learning-based segmentation, classical computer vision approaches, including thresholding and watershed segmentation, were evaluated (Otsu et al., 1979; Vincent and Soille, 1991). These classical methods, while computationally efficient, failed to generalize across all crop species with more complex canopy architectures, thin leaf structures, and nonuniform greenhouse backgrounds, including varying window lighting and neighboring plants. These limitations motivated testing, and ultimately adoption, of deep learning-based segmentation models. The comparative segmentation evaluation included BEN v2 (Li et al., 2024), BiRefNet and BiRefNet Dynamic (Zheng et al., 2024), SAM v2.1 (Kirillov et al., 2023), SAM v3 (Carion et al., 2025), YOLO v11 (Jocher and Qiu, 2024), and YOLO v12 (Tian et al., 2025), along with the classical approaches described above.

Plant detection was performed using YOLO v12 on pseudo-RGB images to identify plant regions and suppress background clutter prior to segmentation. The final configurable pipeline supports BiRefNet and SAM v3 as user-selectable segmentation backbones. BiRefNet applies bilateral feature refinement to preserve thin leaf boundaries while suppressing background artifacts (Zheng et al., 2024), whereas SAM v3 leverages a foundation model architecture with multiscale attention for robust delineation of complex plant structures (Carion et al., 2025). In this pipeline, SAM v3 was applied in a detector-free configuration using a text prompt corresponding to the concept of a plant, enabling direct segmentation of plant regions without bounding-box proposals. Segmentation outputs were mapped back to the four-band multispectral stack to isolate plant pixels for downstream phenotypic analyses. In this software, SAM v3 is the default segmentation option; however, end users can change the segmentation model, such as BiRefNet, through the pipeline configuration. All preprocessing and segmentation code is publicly available in the GitHub repository (https://github.com/Advanced-Vision-and-Learning-Lab/ Plant_Analysis_Tool_Pipeline).

#### 2.4.2 Instance Tracking Across Frames

Instance tracking across the vertically stacked frames, consecutive images captured at successive camera positions along the Z-axis from base of plant to the top of canopy, was performed for each plant using the Segment Anything Model for Long Sequences (SAM2Long) (Ding et al., 2025). SAM2Long extends the original SAM architecture (Kirillov et al., 2023) through temporal attention and a sequence-aware transformer that aligns instance segmentation masks across image sequences. This formulation enables consistent assignment of plant instance identities across frames, even in the presence of overlapping plant structures, self-occlusion, and variations in canopy geometry. Within the pipeline, per-frame segmentation masks generated by user-selected backbones, including SAM v3 (Carion et al., 2025) and BiRefNet (Zheng et al., 2024), were provided as inputs to SAM2Long for cross-frame instance association. SAM v3 was evaluated in standalone mode. By leveraging temporal embeddings and feature matching, SAM2Long produced persistent instance identifiers and reconstructed continuous plant masks across all 13 vertical frames, supporting reliable aggregation of morphological and spectral traits across spatial viewpoints and growth stages.

#### 2.4.3 Image Stitching

Image stitching is essential in greenhouse phenotyping when the plant extends beyond the field of view of a single frame. Tall crops such as mature maize, bioenergy sorghum, or sugarcane exceed the vertical coverage of fixed camera systems, so overlapping views are captured as the camera moves along the stem. Each image contains only a local part of the plant, but stitching them recovers the full architecture and provides a coherent representation for downstream phenotyping tasks. Recent advances in sparse-view 3D reconstruction, particularly Gaussian splatting approaches (Salehi et al., 2026), demonstrate how limited viewpoints can be integrated into geometrically consistent scene representations, highlighting the importance of robust alignment strategies when aggregating partial observations. However, greenhouse imagery presents a challenge for automatic stitching. The background contains dense, repeated structures and neighboring plants that generate many high-contrast keypoints favored by classical feature detectors, while the plant interior offers comparatively fewer stable keypoints beyond leaf boundaries and the central stem. Standard stitching pipelines therefore extract substantially more features from the background than from the plant itself, causing alignment to follow irrelevant structures and introducing geometric distortions into the final mosaic. These challenges motivated the development of plant-aware stitching strategies that prioritize biologically relevant structures while suppressing background-driven alignment artifacts. To address these challenges, image stitching in this pipeline was evaluated using vertically stacked maize plant image sequences and performed using the Scale-Invariant Feature Transform (SIFT) (Lowe, 1999) for keypoint detection. SIFT-based keypoint correspondences between overlapping maize frames were used to estimate a homography transformation, which was applied to warp and align consecutive frames into a single plant mosaic. SIFT was selected due to its robustness and repeatability in detecting stable keypoints across complex maize plant structures, as supported by the quantitative evaluation in Section 3.3.

### 2.5 Feature Extraction Modules

#### 2.5.1 Vegetation Indices

A total of 48 vegetation indices were computed from the segmented multispectral images to quantify plant pigment composition, canopy vigor, and structural traits. Each index was derived from linear or nonlinear relationships among the four spectral bands (green, red, red-edge, and near-infrared), following the general form in Equation 3:

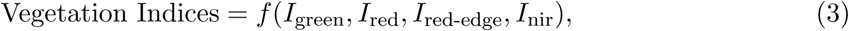

where *I_λ_* represents the reflectance intensity for spectral band *λ*. The indices were calculated using the formulas implemented on segmented plant images. Pixel-wise vegetation index maps were generated, and summary statistics (mean, standard deviation, minimum, and maximum) were computed for each plant instance.

Representative indices included the Normalized Difference Vegetation Index (NDVI) (Rouse Jr et al., 1973), Green NDVI (GNDVI) (Gitelson and Merzlyak, 1998), Normalized Difference RedEdge Index (NDRE) (Barnes et al., 2000), Anthocyanin Reflectance Index (ARI) (Gitelson et al., 2001), Modified Chlorophyll Absorption Ratio Index (MCARI) (Daughtry et al., 2000), and Optimized Soil-Adjusted Vegetation Index (OSAVI) (Rondeaux et al., 1996). Additional indices sensitive to canopy structure, water content, and physiological stress (e.g., MSAVI, TSAVI, CCCI, NDWI) were also implemented (Qi et al., 1994; Baret and Guyot, 1991; Gao, 1995; El-Shikha et al., 2008). All computations incorporated a small numerical constant *ε* = 10*^−^*^6^ to prevent division by zero, consistent with the numerical safeguards used in the processing code. The complete set of 48 vegetation indices, including formulas and spectral dependencies, is provided in Supplemental Table S2.

#### 2.5.2 Morphological Features

Morphological traits were extracted using a hybrid workflow that combined the PlantCV library (Gehan et al., 2017) with supplementary OpenCV-based algorithms (Itseez, 2015). This workflow quantified plant shape, structure, and architectural complexity from binary segmentation masks generated for each image. All trait calculations were restricted to plant pixels as defined by the segmentation outputs to ensure exclusion of background elements. Prior to measurement, binary masks were preprocessed using morphological opening and connected-component filtering to remove background noise and artifacts smaller than 1000 pixels. For each plant, the largest connected contour was selected as the primary region of interest. From this region, basic morphological descriptors were computed, including projected area, perimeter, width, height, bounding-box area, aspect ratio, elongation, circularity, convexity, and solidity. Convex hull analysis was used to estimate canopy compactness, and ellipse fitting was applied to derive major and minor axis lengths for geometric characterization.

Structural topology was further characterized using skeleton-based analysis implemented through the PlantCV morphological pipeline (Gehan et al., 2017). Plant masks were skeletonized and iteratively pruned across multiple scales to suppress minor branches and preserve dominant structural axes. The resulting skeletons were analyzed to identify branch points, tip points, and segmented components corresponding to leaves and stems. From these representations, additional traits such as leaf count, stem count, skeleton length, and branch density were derived. All measurements were converted from pixel units to physical dimensions using a pixel-to-centimeter conversion derived from spatial metadata recorded during image acquisition based on the pinhole camera model (Szeliski, 2022). The working distance *D* between the camera and the plant was estimated from the relative positional metadata (specifically the difference between the Plant Y and Robot Y coordinates, which represents the camera-to-plant distance along the imaging axis). Using known camera parameters (sensor width = 11.26 mm, focal length = 6 mm, image width = 512 px), the field of view was computed as FOV_width_ = D・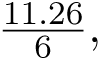 and the pixel-to-centimeter scale was obtained as 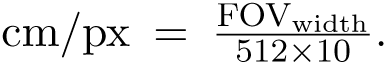 This approach removes the requirement for a physical calibration marker, as the necessary spatial metadata are automatically recorded for each image acquisition. Extracted morphological traits were archived together with diagnostic visualizations, including plant contours, skeleton overlays, and keypoint maps, to support quality assurance, reproducibility, and downstream phenotypic analysis.

### Self-supervised keypoint detection

To enhance the robustness of morphological feature extraction without relying on labor-intensive manual keypoint annotation of greenhouse images, a self-supervised learning (SSL) framework was employed for the automatic detection of structural keypoints such as leaf tips. Self-Distillation with No Labels (DINO) (Caron et al., 2021), a self-supervised vision transformer framework, was used as the pretext task to train a model using the curated PGP v2 dataset, consisting of 1,378 unlabeled pseudo-RGB images of sorghum plants. The PGP v2 unlabeled images were used exclusively for DINO pretraining and were not combined with the Sorghum Leaf Counting Dataset (Miao et al., 2021), described below, used for supervised finetuning; these two datasets were kept entirely separate throughout all stages of training and evaluation. During this pretraining stage, the model learned structure-aware representations by aligning image patches across diverse instances of sorghum plants with varying genotypes, growth stages, and architectural patterns. This process emphasized spectral–spatial continuity, shape geometry, and fine structural cues without the use of manual labels.

The pretrained encoder was subsequently finetuned for keypoint detection using a subset of the publicly available Sorghum Leaf Counting Dataset (Miao et al., 2021), enabling supervised learning of leaf tip–related structural cues without requiring manual keypoint annotation of greenhouse images. This dataset contains 27,770 cropped RGB images of sorghum plants acquired under controlled greenhouse conditions (different from APPG), accompanied by annotations specifying image filename, genotype identifier, total leaf count, and viewing angle. The dataset was designed for automated leaf counting and structural trait analysis in grain crops. Accordingly, the detected structural keypoints were used to support leaf tip localization and downstream leaf counting tasks. For this stage, a detection pipeline based on the You Only Look Once version 12 (YOLOv12) architecture (Tian et al., 2025) was employed, which integrates attention-based mechanisms for real-time object detection. Finetuning was performed using 896 × 896 pixel images, the AdamW optimizer (Loshchilov and Hutter, 2017), and a cosine annealing learning rate schedule (Loshchilov and Hutter, 2016) over 100 epochs. The Sorghum Leaf Counting Dataset was divided into 60% for training, 20% for validation, and 20% for independent testing, with no overlap between splits. The independent test set was held out entirely during model development and used solely for final performance evaluation. All model development was executed on the Texas A&M High Performance Research Computing (HPRC) cluster using compute nodes equipped with dual NVIDIA A100 (40 GB) GPUs.

#### 2.5.3 Texture Features

Texture features were extracted to quantify microstructural and spatial heterogeneity in plant surfaces, which can reflect physiological variation such as leaf venation density, surface roughness, and canopy organization. The pipeline implemented a comprehensive texture analysis framework comprising four complementary descriptors: Local Binary Pattern (LBP), Histogram of Oriented Gradients (HOG), Lacunarity (including a Differential Box Counting variant), and Edge Histogram Descriptor (EHD). Each descriptor captures distinct aspects of texture geometry, contrast, and orientation. The LBP operator (Ojala et al., 2002) encodes local contrast by thresholding neighborhood intensities around each pixel, producing rotationand illuminationinvariant maps of fine-scale surface texture. HOG (Dalal and Triggs, 2005) captures macroscopic structural patterns by aggregating local edge orientation histograms computed over overlapping spatial cells, emphasizing directional gradients associated with leaf edges and venation patterns. Lacunarity features were computed to quantify textural heterogeneity and the distribution of spatial gaps (Plotnick et al., 1993; Allain and Cloitre, 1991). Three lacunarity types were implemented: (i) local single-window lacunarity, which estimates the variance-to-mean ratio of pixel intensities within a fixed sliding window (Tolle et al., 2003); (ii) multi-scale averaged lacunarity, which aggregates measurements across multiple kernel sizes to capture hierarchical texture variation (Dong et al., 2017); and (iii) Differential Box Counting (DBC) Lacunarity (Mohan and Peeples, 2024), implemented as a PyTorch-based model following the DBC formulation for fractal texture estimation. The DBC model computes texture irregularity by applying local max–min pooling over sliding windows. The Edge Histogram Descriptor (EHD) (Manjunath et al., 2001) quantifies the spatial distribution of edge directions by convolving images with edge filters rotated at 45*^◦^* intervals between 0 and 315*^◦^* (inclusive). The filters capture horizontal, diagonal, vertical, and anti-diagonal edges, while a threshold is used to capture low responses (i.e., “no edge”). The maximum directional edge responses were aggregated through histograms that describe structural anisotropy and leaf alignment.

Texture features were extracted across multiple imaging domains to capture both spectral and structural diversity. Specifically, six image modalities were analyzed per plant instance: pseudo-color composite, NIR, red-edge, red, green, and a principal component analysis (PCA) representation. The pseudo-color composite (originally a 3-channel RGB image) was converted to grayscale using the cvtColor function in OpenCV (Itseez, 2015) before texture feature extraction. To generate the PCA representation, the four multispectral bands were first stacked into a per-pixel feature vector and then masked to plant regions (pixels inside the plant masks). Here, *H* and *W* denote the image height and width in pixels, respectively. Pixels outside the plant mask were set to NaN (not a number) and excluded from PCA. The stacked image cube of size *H* × *W* × 4 was reshaped into a (*HW*) × 4 matrix, and rows containing NaNs were removed. PCA was then fit using scikit-learn (Pedregosa et al., 2011) with one component and whitening enabled, after which the first principal component score was computed for each valid pixel. The resulting *HW*-length score vector was reshaped back to an *H* × *W* image. Within the plant mask, this PCA score image was min–max normalized and converted to an 8-bit grayscale image for downstream texture computation. Each grayscale band image was processed using the full texture feature suite, and all computations were masked to plant regions to exclude background noise. For each descriptor and imaging domain, pixel-level feature maps were summarized by statistical aggregation measures (mean, standard deviation, minimum, maximum, and median) to generate interpretable quantitative profiles of plant surface texture.

### 2.6 Temporal and Statistical Analysis

All features (i.e., vegetation indices, morphological, and texture) extracted from segmented plant regions were aggregated by plant identity and imaging date to construct temporal feature matrices. Plant identity across images was established using the SAM2Long temporal tracking framework described in Section 2.4.2, which assigns a persistent instance identifier to each plant by leveraging spatial consistency and mask overlap across consecutive frames. This enables the same plant to be tracked across vertically stacked images and across imaging sessions. For each spectral band (green, red, red-edge, and near-infrared) and corresponding derived component (e.g., PCA projections and vegetation indices), statistical descriptors were computed to summarize the pixel-level reflectance distributions. These descriptors included the mean, standard deviation, minimum, maximum, median, interquartile range (difference between the 25th and 75th percentiles), skewness, kurtosis, and Shannon entropy. Together, these metrics capture both central tendency and higher-order variability in spectral responses within plant tissues. Temporal aggregation across imaging sessions provided continuous profiles of each plant’s reflectance and morphological properties, enabling visualization of dynamic patterns in growth and canopy physiology. The resulting feature matrices form the basis for downstream analyses and longitudinal comparison of temporal trajectories across species, treatments, or developmental stages. To evaluate treatment-level differences, statistical analysis was performed using a linear mixed-effects model (LMM) (Lindstrom and Bates, 1990) incorporating repeated measurements over time. In addition, control-mean t-tests and robust z-tests (Huber, 1981) were applied across vegetation index features, with multiple hypothesis testing controlled using the Benjamini–Hochberg false discovery rate (FDR) (Benjamini and Hochberg, 1995) correction.

### 2.7 Case Study Experimental Details

For the sorghum treatment-level case study, mutagenized sorghum lines were generated by applying Low Energy Electron Beam (LEEB) irradiation to sorghum seeds under different treatment parameters using the EBLab200 SKAN Stein AG low-energy eBeam at the TAMU eBeam Facility. A total of 48 M_2_ generation plants from the sorghum background genotype TAMU breeding line R.08306 were evaluated. The experiment included seven LEEB treatment groups (G1–G7) and one non-treated control group (NT, Group 8), with six plants per group in the APPG. The objective was to evaluate whether the pipeline could detect subtle phenotypic differences induced by different LEEB treatment parameters (beyond what is visually observable by researchers). For each plant and imaging date, vegetation-index-derived features were extracted from segmented plant regions. The pipeline initially computed 384 features from 48 multispectral vegetation indices and eight summary statistics per index: mean, standard deviation, minimum, maximum, median, 25th percentile, 75th percentile, and undefined-pixel fraction. These were reduced to 240 vegetation-index features by removing the two per-index extrema (minimum and maximum), the per-index undefined-pixel fraction, and any remaining zero-variance features; no feature exceeded the 50% missing-value threshold.

Two complementary statistical analyses were performed. First, individual plants were ranked by divergence from the non-treated control using temporal mean and time-normalized area-under-the-curve summaries for each retained feature. Control-mean *t*-tests and robust median/MAD-based *z*-tests were computed for each plant and feature, with Benjamini–Hochberg false discovery rate correction applied across the 240 features for each plant. Second, group-level treatment effects were evaluated separately for each retained feature using a Linear MixedEffects Model (LMM) (Lindstrom and Bates, 1990). For each feature, the response variable was the vegetation-index feature value, treatment group and linear imaging day were included as fixed effects, and plant identity was included as a random intercept to account for repeated measurements over time. The covariance structure was defined by the plant-level random intercept, with independent residual errors. Models were fit using the mixedlm implementation of MixedLM from the Python statsmodels package (Seabold et al., 2010). Treatment-versus-control contrasts were obtained from the treatment-group fixed-effect coefficients, and *p*-values were adjusted within each treatment group across the 240 features using the Benjamini-Hochberg false discovery rate correction.

To evaluate the generalizability of the pipeline beyond the APPG multispectral greenhouse system, the framework was applied to a maize cold stress experiment from the CERCA (Circular Economy that Reimagines Corn Agriculture) project at the University of Nebraska–Lincoln. Ten maize genotypes, with one plant per genotype, were grown in Pro-Mix BX in 5.1 cm diameter peat pots in a growth chamber for two weeks at 29*^◦^*C/27*^◦^*C under a 16 h day/8 h night cycle to allow germination and establishment. Plants were then transferred to chambers applying one of three temperature treatments under a 12 h day/12 h night cycle: (1) 20*^◦^*C for two weeks, (2) 10*^◦^*C for one week, or (3) 4*^◦^*C for three days. Transfer was always conducted at the start of the respective night period. After each treatment, plants were allowed to recover under greenhouse conditions until the V7 growth stage. Light conditions were approximately 300 *µ*mol photons m*^−^*^2^ s*^−^*^1^ in the chambers. The ten maize hybrid lines were LH287 × LH244, CG119 × CG108, PH09B × PH2N0, HC33 × PH1B5, PHR03 × LH244, PH09B × PH2MW, PHB47 × PHZ51, PHJ40 × PHAJ0, B73 × Mo17, and Tx777 × LH195. Images were acquired immediately prior to treatment, within a three-hour window immediately following treatment, and once the V7 stage was reached during recovery using a Canon EOS M10 RGB camera.

For each maize plant image, the pipeline generated a structured feature table containing 620 numerical phenotypic features. These features included RGB-based vegetation index statistics from 11 indices, including ExG, ExGR, MGRVI, NGRDI, and VARI; morphological descriptors, including size, skeleton structure, curvature, branch points, and tip detection; and texture descriptors extracted across five imaging representations, including blue, green, red, color composite, and PCA-derived representations. The resulting feature matrix was used for downstream multivariate analysis. Principal component analysis (PCA) (Abdi and Williams, 2010) was then applied to the normalized RGB-derived vegetation index features to evaluate whether the extracted image features captured variation across the cold stress timeline.

## 3 Results and Discussion

### 3.1 Plant Segmentation Performance

The analytical pipeline performs automated plant segmentation to separate plant from background across greenhouse images including multiple crop species and growth stages. In most experiments, before segmentation can occur, the plants must be detected (i.e., object detection). Based on the comparative evaluation described in Section 2.4.1, this section reports the qualitative and quantitative segmentation performance of the evaluated methods.

Qualitative plant segmentation results across six representative samples, including sorghum, maize, and cotton (Figure 4), highlighted performance differences across models and species. BEN v2 successfully removed background clutter but occasionally over-segmented regions with complex plant geometry. BiRefNet and BiRefNet Dynamic produced high-quality masks, particularly in dense canopies with overlapping foliage, and yielded identical results in our experiments. SAM v2.1 showed limited performance for these greenhouse images. SAM v3 produced the most accurate segmentation masks overall, showing clear alignment with plant boundaries and reliable preservation of thin and curved leaf structures. YOLO v11 and YOLO v12 generated reliable plant masks, although their boundary detail was generally lower than that of the stronger segmentation models.

**Figure 4:**
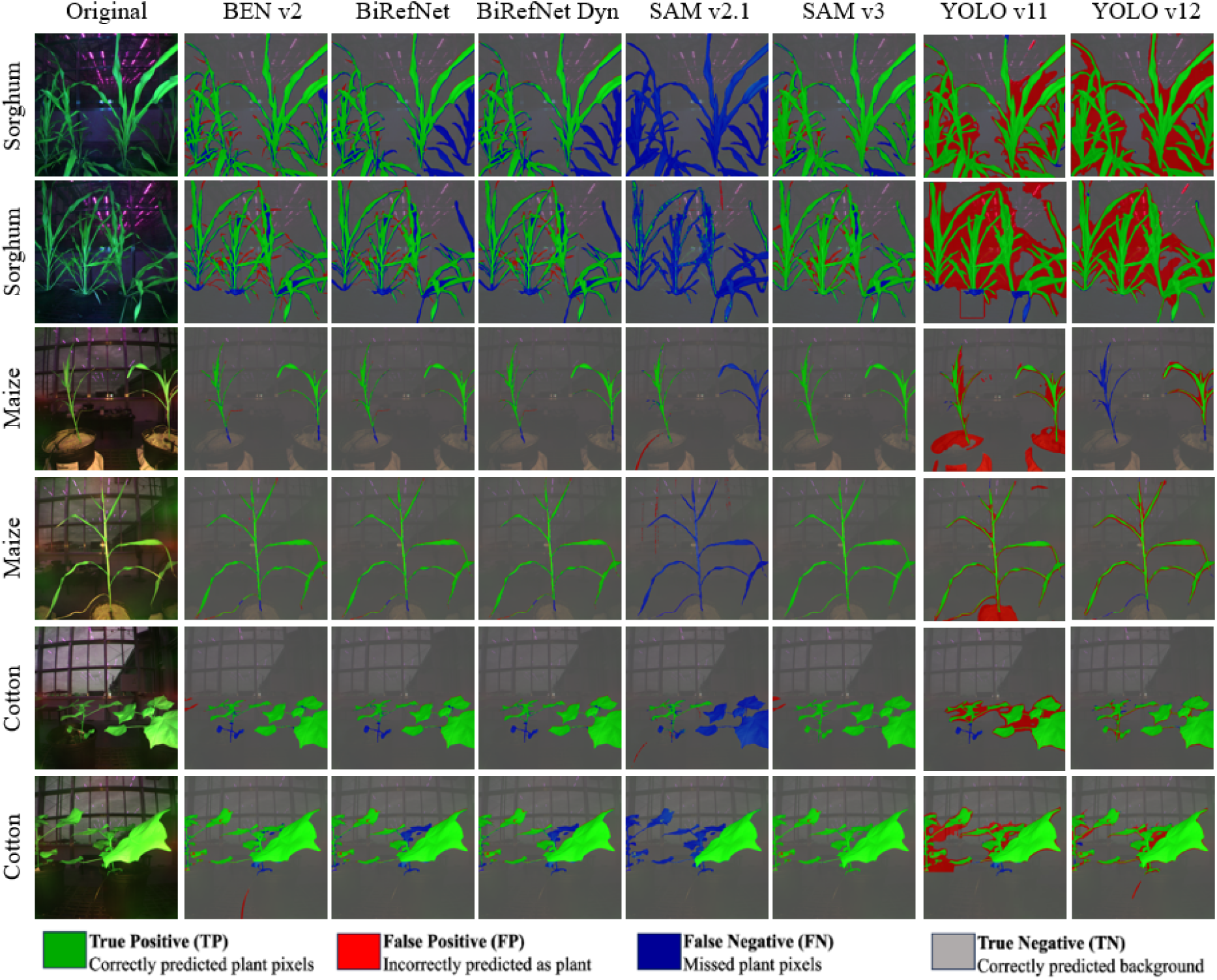
Qualitative comparison of plant segmentation results across multiple models and crop species.

In addition to qualitative comparisons, Table 3 reports quantitative segmentation accuracy on human-annotated masks. Consistent with the qualitative results, SAM v3 achieved the highest IoU and Dice scores, followed by BiRefNet-based models, while YOLO-based approaches showed competitive but lower boundary-level accuracy. Overall, detection-guided segmentation improved robustness to background variation and pot-related artifacts, whereas detector-free segmentation with SAM v3 provided superior preservation of thin plant structures, supporting reliable downstream phenotypic analysis.

To further contextualize the performance gains of deep learning-based segmentation, a qualitative comparison between classical computer vision approaches (Otsu thresholding and watershed segmentation) and SAM v3 across the four crop species is provided in Supplemental Figure S1. Table 2 summarizes the corresponding IoU scores and processing times. Processing times for these classical methods were measured on an Intel i9-10920X CPU, whereas SAM v3 inference was performed on an NVIDIA RTX 4090 GPU. The choice of CPU or GPU was dictated by the requirements of each algorithm. Classical methods were substantially faster but produced lower and more variable segmentation accuracy across species (Table 2), whereas SAM v3 consistently achieved stronger IoU performance with lower variability. These results indicate that the accuracy gains of deep learning-based segmentation justify the additional computational cost, particularly for species with complex canopy architectures such as sorghum and rice, where classical methods failed most severely.

**Table 2:**
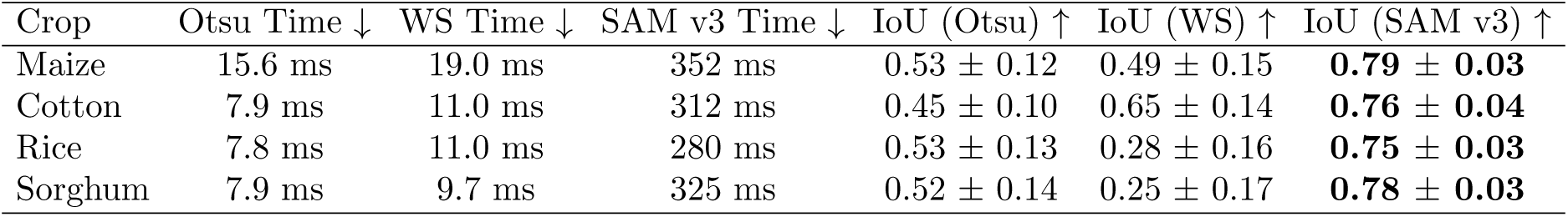
Per-crop comparison of classical segmentation methods and SAM v3 in terms of computational efficiency (inference time) and segmentation accuracy (IoU). Processing times are reported in milliseconds and represent the average inference time per image across all plants for each crop type. IoU values represent mean ± standard deviation across samples for each crop type. WS (watershed segmentation); SAM (Segment Anything Model)

**Table 3:** Quantitative comparison of segmentation accuracy across models using all annotated plant ground-truth masks from the evaluation dataset. Predicted plant masks were compared with manual ground-truth masks, and values are reported as mean ± standard deviation across the evaluation set. SAM v3 achieved the highest overall segmentation performance.

| Model | IoU ↑ | Dice ↑ | Precision ↑ | Recall ↑ |
| --- | --- | --- | --- | --- |
| BEN v2 | $0.63 \pm 0.11$ | $0.77 \pm 0.09$ | $0.90 \pm 0.04$ | $0.69 \pm 0.13$ |
| BiRefNet | $0.72 \pm 0.17$ | $0.81 \pm 0.16$ | $0.86 \pm 0.07$ | $0.77 \pm 0.09$ |
| BiRefNet Dynamic | $0.72 \pm 0.17$ | $0.81 \pm 0.16$ | $0.86 \pm 0.07$ | $0.77 \pm 0.09$ |
| SAM v2.1 | $0.18 \pm 0.12$ | $0.29 \pm 0.18$ | $0.76 \pm 0.04$ | $0.18 \pm 0.12$ |
| SAM v3 | **$0.78 \pm 0.04$** | **$0.88 \pm 0.02$** | **$0.92 \pm 0.04$** | $0.84 \pm 0.04$ |
| YOLO v11 | $0.43 \pm 0.12$ | $0.58 \pm 0.11$ | $0.47 \pm 0.15$ | $0.84 \pm 0.05$ |
| YOLO v12 | $0.59 \pm 0.06$ | $0.74 \pm 0.05$ | $0.61 \pm 0.07$ | **$0.96 \pm 0.03$** |

### 3.2 Temporal Instance-Level Tracking Performance

Temporal instance-level tracking was evaluated qualitatively to assess the ability of different segmentation configurations to preserve plant identities across multi-frame greenhouse sequences.

The evaluation focused on visually diagnosing identity stability, instance fragmentation, merging, and missed detections across vertically stacked frames acquired along the plant stem. Two configurations were compared: SAM v3 used in standalone mode with internal identity propagation, and BiRefNet combined with SAM2Long, where per-frame segmentation masks are explicitly associated across frames using a sequence-aware tracker.

As the greenhouse camera captures images of tall plants from different spatial positions across frames, it is important that plants are detected and segmented consistently in each frame and that the identity of each plant is preserved. Figure 5 illustrates representative results on sorghum (7 frames) and maize (5 frames) sequences. In the sorghum sequence, BiRefNet and SAM2Long consistently maintain the same instance identities across frames, whereas SAM v3 exhibits an identity switch between mid-sequence frames 3 and 4 for the same plant. In the maize sequence, BiRefNet and SAM2Long occasionally merge instances or miss a plant under limited spatial separation, while SAM v3 alternates between over-segmentation, identity fragmentation, and instance merging. These results highlight that explicit temporal association improves identity stability but remains sensitive to dense canopies and overlapping plant structures. To overcome the issue of plant identity swapping, we determined that image stitching between frames was needed.

**Figure 5:**
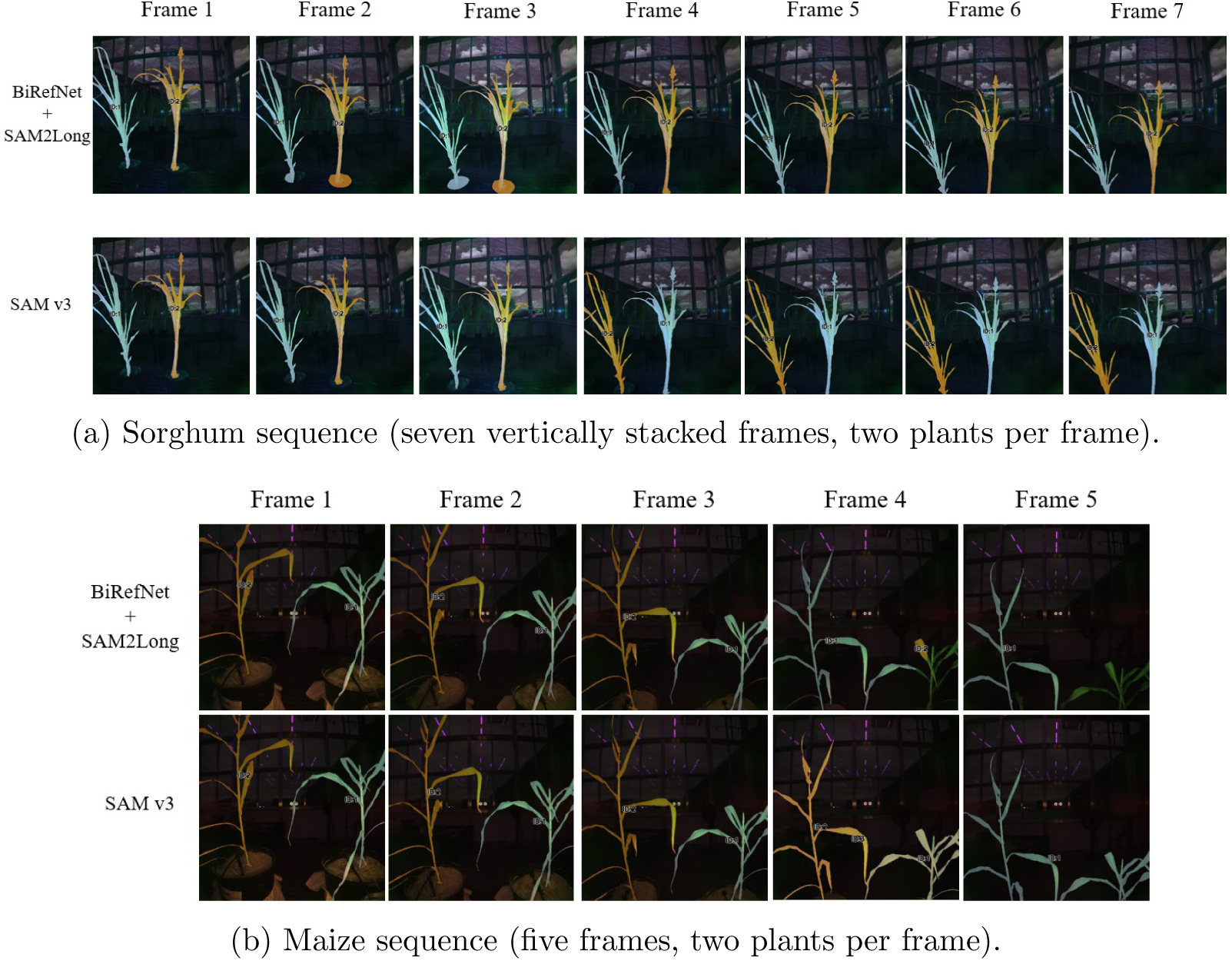
Qualitative comparison of unsupervised temporal instance-level tracking. For each subplot, the top row shows results from BiRefNet and SAM2Long, and the bottom row shows SAM v3. Colors and numeric labels denote plant instance identities. BiRefNet and SAM2Long generally preserve identities more consistently, while SAM v3 exhibits identity swaps, oversegmentation, and identity merging under challenging spatial configurations.

**Figure 6:**
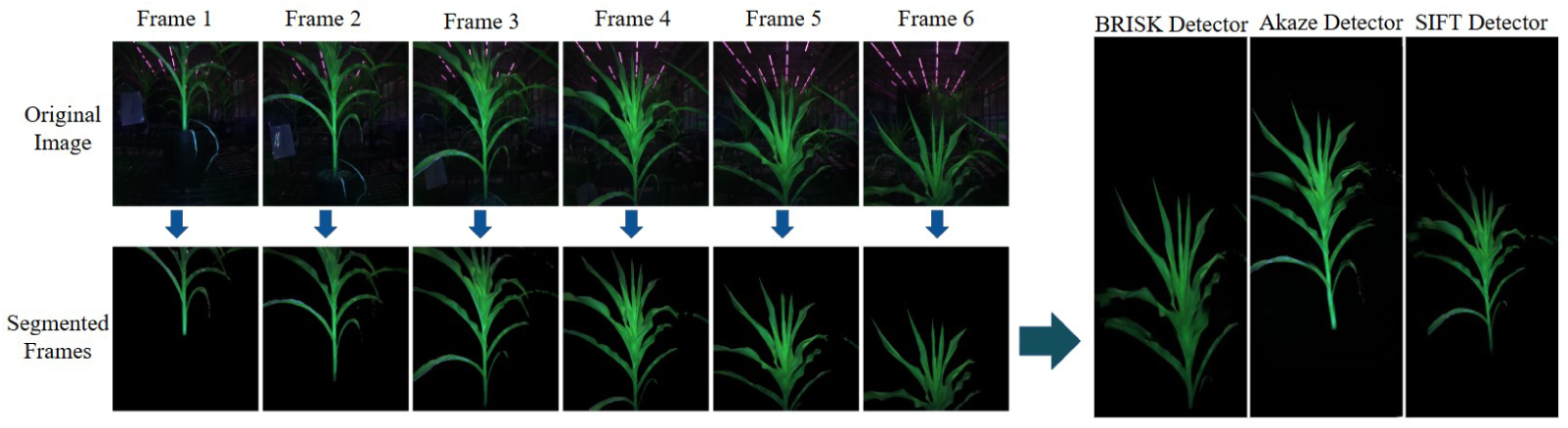
Qualitative stitching results for a maize plant. Six consecutive original frames (top) and their corresponding SAM v3 segmented frames (bottom) were used as inputs to the stitching pipeline. The right side presents stitched mosaics obtained using three different feature detectors (BRISK, AKAZE, and SIFT), illustrating the impact of feature detector selection on reconstruction quality in the absence of a single full-plant panorama.

### 3.3 Image Stitching Performance

Keypoint (also called tiepoint) detection was required for image stitching because reliable tiepoint correspondences between overlapping frames were needed to estimate the geometric transformation (homography) and align frames into a single mosaic. Stitching was evaluated using three keypoint detectors: Binary Robust Invariant Scalable Keypoints (BRISK) (Leutenegger et al., 2011), Accelerated-KAZE (AKAZE) (Alcantarilla and Solutions, 2011), and the Scale-Invariant Feature Transform (SIFT) (Lowe, 1999). Although BRISK and AKAZE produced acceptable alignments in some cases, they yielded fewer reliable matched correspondences than SIFT for the representative sequence evaluated. This limitation can arise from repetitive leaf patterns and low-texture regions common in plant imagery, which may lead to incomplete mosaics or skipped frame-to-frame alignments. In contrast, SIFT produced the highest average number of matched correspondences and successfully linked all available frame-to-frame matching opportunities, supporting more stable mosaic reconstruction.

Since a panoramic reference was generally not available, stitching performance was assessed through qualitative visual inspection of alignment consistency across vertically overlapping frames. The technical efficiency of the keypoint detectors during the reconstruction process was summarized in Table 4. The values reported in this table correspond to one representative maize plant sequence with 12 available frame-to-frame matching opportunities. While the keypoints column represents the total number of features detected in this representative sequence, the most critical factor for stitching was how many of these features matched within the overlapping regions of consecutive frames. Although AKAZE detected the largest number of keypoints, SIFT produced the highest average number of successful matches between consecutive frames. The successful pairs column indicates the number of consecutive image pairs, out of the 12 available frame-to-frame matching opportunities for this maize sequence, which were effectively linked and added to the final mosaic. This metric shows how many frame-to-frame alignments were retained by the algorithm and how many were automatically discarded because a suitable match could not be found. BRISK linked only 10 of the 12 available matching opportunities, meaning that two frame-to-frame alignments were excluded due to poor alignment, whereas both AKAZE and SIFT linked all 12. Overall, SIFT provided the most reliable stitching performance because it achieved the highest average number of feature matches while also successfully linking all available matching opportunities.

**Table 4:**
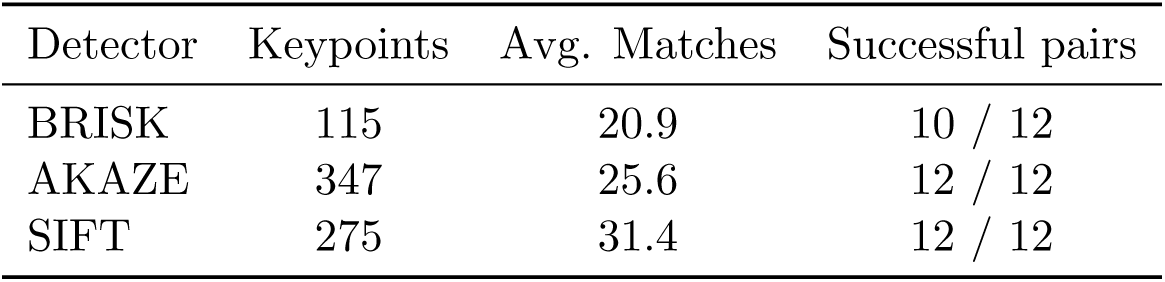
Feature detector statistics for stitched construction. The table reports the number of detected keypoints, the average number of feature matches per source image, and the number of consecutive image pairs that successfully yielded a valid homography.

| Detector | Keypoints | Avg. Matches | Successful pairs |
| --- | --- | --- | --- |
| BRISK | 115 | 20.9 | 10 / 12 |
| AKAZE | 347 | 25.6 | 12 / 12 |
| SIFT | 275 | 31.4 | 12 / 12 |

### 3.4 Plant Tip Detection Performance

An SSL framework based on DINO was employed to improve structural keypoint detection within the greenhouse phenotyping pipeline. The model was initially trained on unlabeled pseudo-RGB PGP sorghum images to learn spatially aware representations through a distillation-based pretext task and subsequently fine-tuned for supervised keypoint detection using labeled data. To assess the impact of SSL pretraining, the SSL-based approach was evaluated against two baselines: (1) a conventional supervised detector trained for bounding-box localization only, and (2) a fully supervised pose estimation model trained without SSL initialization. The supervised baseline refers to a YOLOv12 pose estimation model trained from scratch using random weight initialization and no self-supervised pretraining, allowing the effect of DINO-based SSL pretraining to be isolated. Evaluation followed standard supervised YOLOv12 pose metrics, including Precision (P), Recall (R), mean Average Precision at 0.50 IoU (mAP_50_), and mean Average Precision averaged across IoUs from 0.50 to 0.95 (mAP_50-95_). Metrics were computed separately for bounding-box detection (“Box”) and keypoint estimation (“Pose”).

The SSL-initialized model achieved higher accuracy in keypoint detection than both supervised baselines (Table 5). The model reached a precision of 84%, recall of 85%, mAP_50_ of 0.90, and mAP_50-95_ of 0.89, compared to an mAP_50-95_ of 0.50 for the fully supervised pose estimation model trained without SSL pretraining. Bounding-box detection remained more challenging across all methods, with the SSL-enhanced model achieving an mAP_50-95_ of 0.27, likely due to ambiguous canopy boundaries and overlapping leaf structures. These results indicate that SSL pretraining improves the transferability of learned structural representations for plant tip detection under varying canopy architectures and illumination conditions, while reducing reliance on manually annotated training data. The SSL-based training and transfer pipeline highlighted how DINO pretraining on unlabeled greenhouse images improves downstream plant tip keypoint detection (Figure 7).

**Figure 7:**
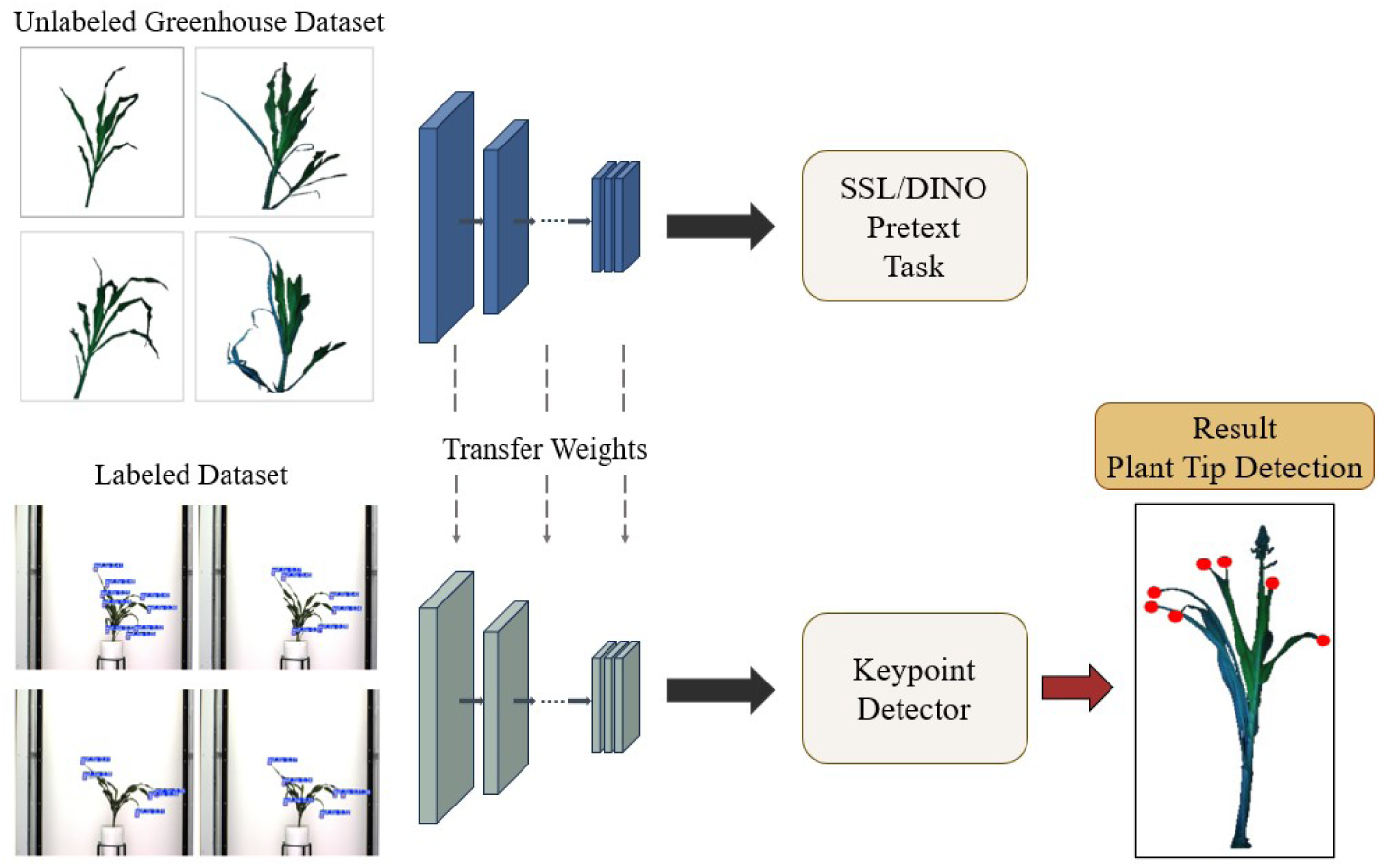
Overview of the self-supervised plant tip detection framework. A DINO-based selfsupervised model is first pretrained on unlabeled pseudo-RGB greenhouse images using a distillation pretext task. The learned representations are then transferred and fine-tuned on a labeled dataset for supervised keypoint detection, resulting in accurate plant tip localization.

**Table 5:** Comparison of keypoint detection accuracy for different methods. “Baseline: supervised” refers to a standard supervised YOLOv12 pose estimation model trained from scratch using random weight initialization and no self-supervised pretraining, serving as a reference to isolate the contribution of DINO-based SSL pretraining.

| Method | Task | Precision (P) | Recall (R) | $mAP_{50}$ | $mAP_{50-95}$ |
| --- | --- | --- | --- | --- | --- |
| Baseline: supervised | Box | 0.45 | 0.43 | 0.42 | 0.21 |
| Baseline: supervised | Pose | 0.71 | 0.72 | 0.75 | 0.50 |
| DINO + fine-tuning | Box | 0.68 | 0.67 | 0.66 | 0.27 |
| DINO + fine-tuning | Pose | **0.84** | **0.85** | **0.90** | **0.89** |

**Table 6:**
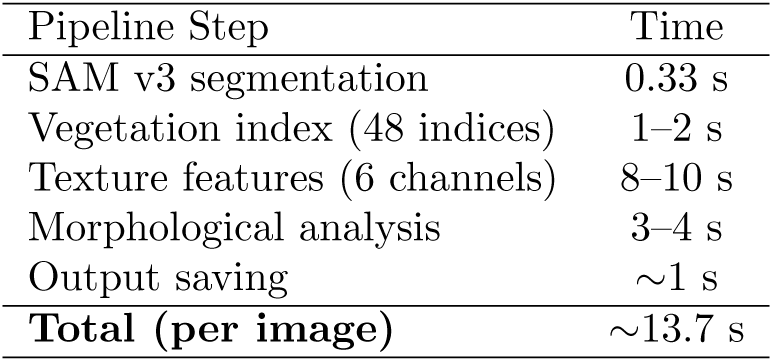
Per-step processing time for one stitched plant image. Times exclude the 3.3-s modelloading overhead and are reported in seconds per image.

### 3.5 Temporal Dynamics of Extracted Features

For each plant and imaging date, a total of 863 quantitative features were extracted (Supplemental Table S3), spanning vegetation indices, spectral statistics, texture descriptors, and morphological traits. This high-dimensional feature set extends beyond conventional phenotyping traits and provides a broad quantitative description of plant spectral, textural, and structural variation. While some extracted features are commonly used in plant phenotyping, such as NDVI, many others provide novel information for future experiment-specific biological discovery. The pipeline additionally enables longitudinal tracking of individual features for each plant across imaging dates. As an example of an interpretable phenotype, NDVI was tracked over time for a representative sorghum plant, illustrating how temporal feature trajectories can capture changes in plant reflectance during growth and development (Figure 8a).

**Figure 8:**
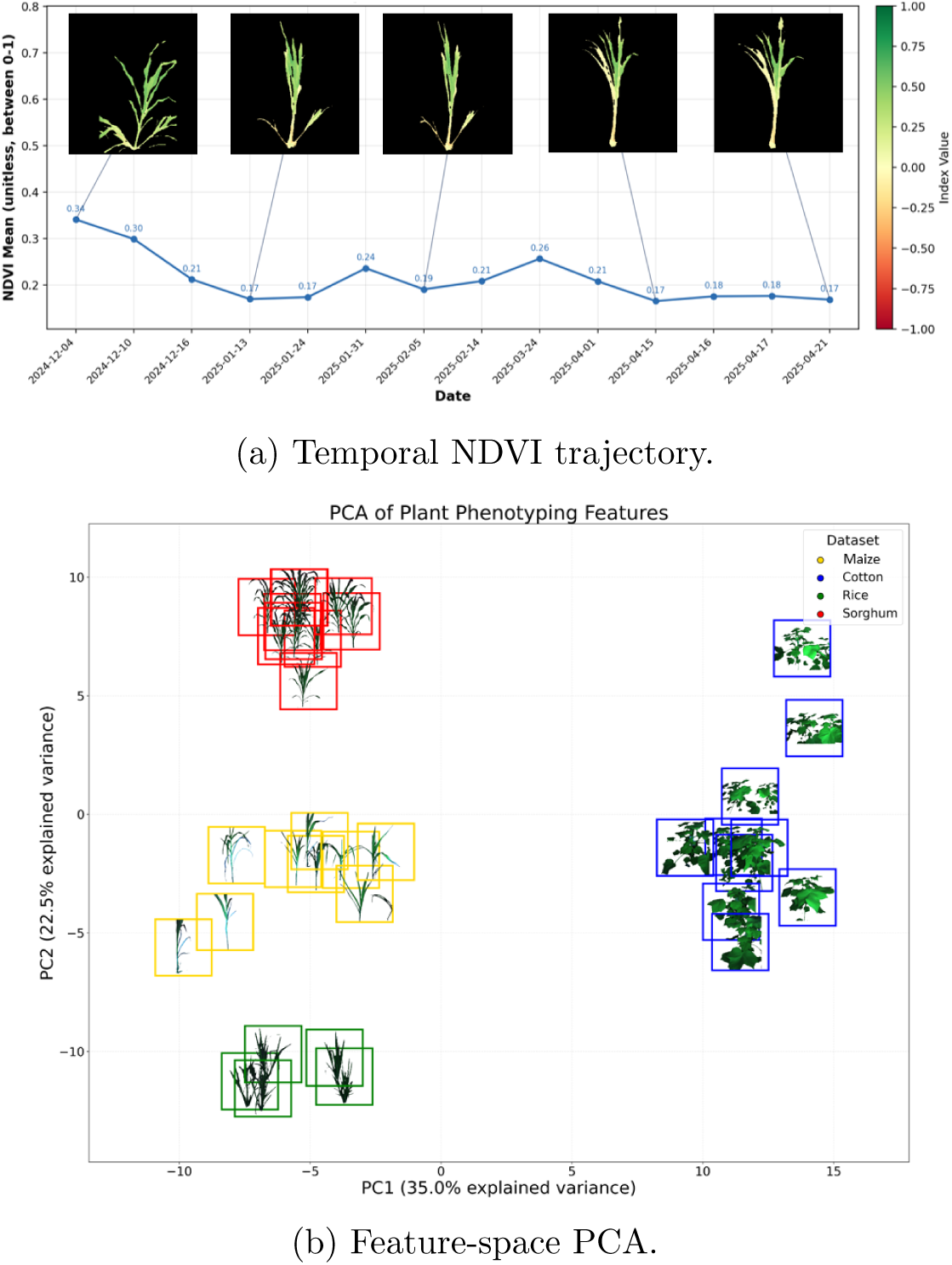
Temporal tracking and high-dimensional feature-space analysis of extracted phenotypic traits. (a) Representative NDVI mean trajectory over time for a single sorghum plant across 17 imaging days from Dec 2024 to Apr 2025. NDVI is shown as an example phenotype to illustrate temporal feature tracking across imaging dates, with five representative images showing plant appearance at selected dates. (b) PCA of the full 863-dimensional phenotyping feature set across 36 crop samples, including maize (*n* = 9), cotton (*n* = 10), rice (*n* = 7), and sorghum (*n* = 10).

To examine the global structure of the extracted feature space, principal component analysis (PCA) (Abdi and Williams, 2010) was performed on the full 863-dimensional phenotyping feature set across all crop samples shown in Figure 8b, including maize, cotton, rice, and sorghum from the PGP v2 dataset. Together, these features captured complementary aspects of plant structure and physiology and enabled multi-dimensional temporal characterization. Temporal profiling from December 2024 to April 2025 revealed coordinated variation across spectral, textural, and morphological domains, reflecting developmental progression and canopy maturation. Because the full 863-feature temporal profile is too complex to present feature by feature, the NDVI trajectory is shown only as a representative example of longitudinal feature tracking, while PCA summarizes broader variation across the high-dimensional feature space across crops.

### 3.6 Texture Feature Extraction

Texture features (Peeples et al., 2021), as extracted in this pipeline, have been less commonly used in controlled-environment plant phenotyping than conventional spectral indices and morphological traits, but can capture complementary information related to leaf surface structure, canopy organization, and spatial heterogeneity. As a case study example, representative texture feature maps were extracted from a cotton plant image using the pseudo-RGB composite converted to grayscale, as described in Section 2.5.3. Four complementary descriptors were computed (Figure 9): Local Binary Pattern (LBP), Histogram of Oriented Gradients (HOG), lacunarity, and Edge Histogram Descriptor (EHD). LBP encoded fine-scale local contrast patterns corresponding to leaf surface microstructure. HOG highlighted macroscopic structural gradients associated with leaf edges and venation. Lacunarity captured textural heterogeneity and the spatial distribution of gaps within the canopy. EHD quantified the directional distribution of edges, reflecting leaf orientation and structural anisotropy. Together, these textural descriptors provide a rich characterization of plant surface properties that complements the spectral and morphological features extracted by the pipeline. These features are further summarized through statistical descriptors and integrated into downstream analyses for phenotypic characterization and comparison across plants and conditions.

**Figure 9:**
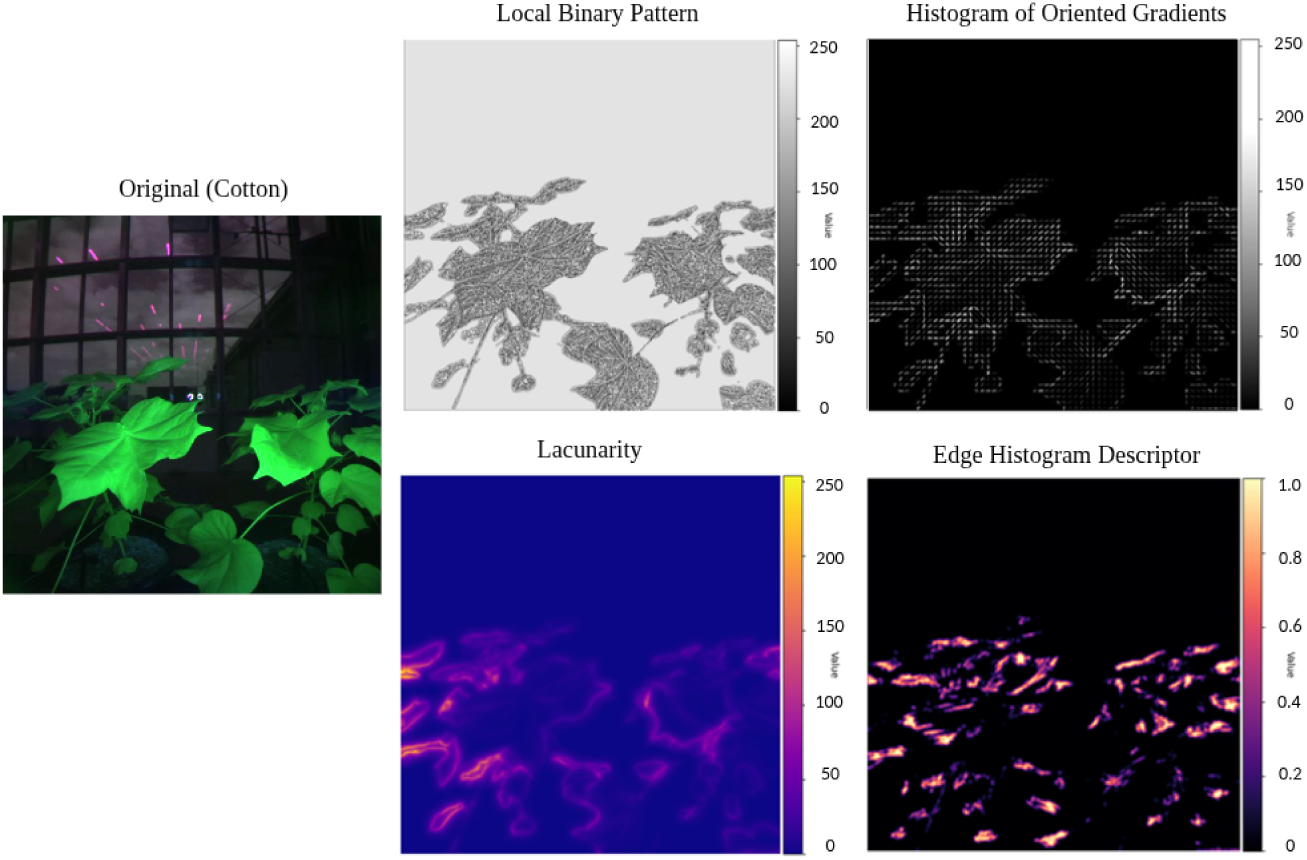
Representative texture feature maps (LBP, HOG, lacunarity, and EHD) extracted from a cotton plant image using the pseudo-RGB composite converted to grayscale, illustrating complementary structural characteristics captured by the pipeline.

**Figure 10:**
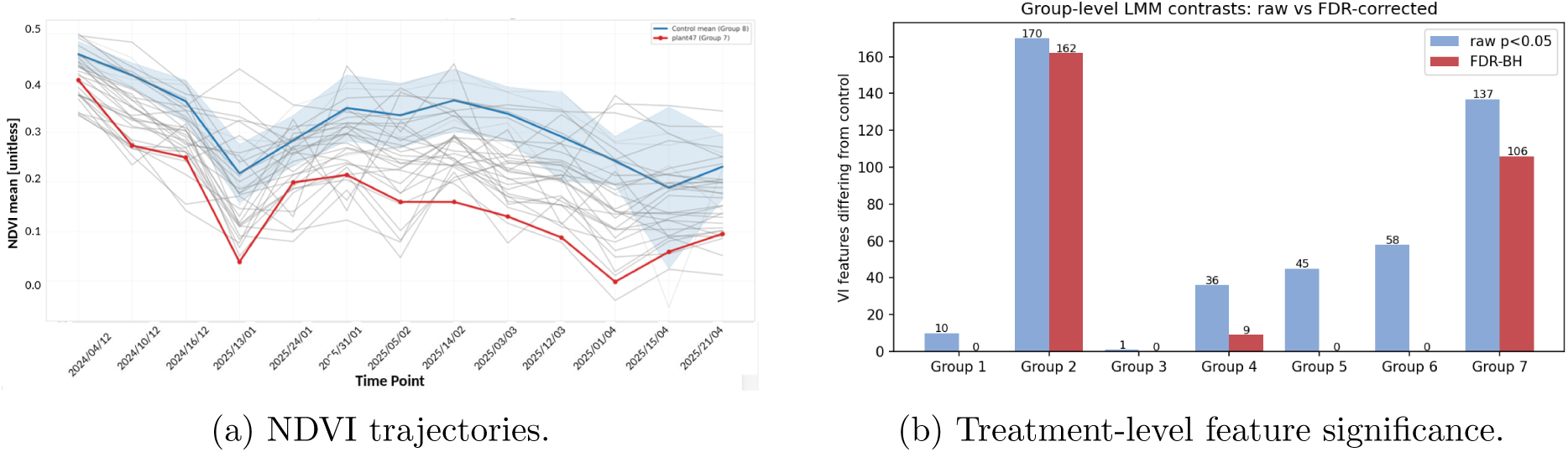
Statistical analysis for the mutagenized sorghum case study. (a) NDVI mean trajectories for treated plants and the non-treated control, with Plant47 (Group 7) highlighted in red. (b) Group-level linear mixed-effects analysis comparing each treatment group with the non-treated control across 240 vegetation-index features. Bars show raw-significant features (*p <* 0.05) and Benjamini–Hochberg FDR-significant features.

### 3.7 Pipeline Efficiency and Scalability

The full pipeline was benchmarked on an NVIDIA GeForce RTX 4090 GPU (24 GB VRAM) paired with an Intel i9-10920X CPU (24 cores). Table 6 reports the per-step processing times for a single stitched image after model loading (cold-start model loading: 3.3 s). Texture feature extraction is the primary computational bottleneck, accounting for approximately 50–60% of total processing time.

For each stitched plant image, processing time was measured on the final stitched image used for feature extraction rather than on individual image frames. Based on an average processing time of approximately 13.7 s per stitched image, the sorghum longitudinal dataset (48 plants × 17 imaging dates; 816 stitched images total) required approximately 3.1 hours for feature extraction. The pipeline outputs a ready-to-use 863-dimensional comma-separated value (CSV) feature matrix with no manual intervention beyond initial configuration.

### 3.8 Case Study

#### 3.8.1 Case Study 1: Treatment-Level Comparison

Case Study 1 on sorghum evaluated whether the pipeline could detect subtle phenotypic responses among sorghum plants subjected to different LEEB mutagenesis treatment parameters (treatment groups 1–7 and a non-treated control; experimental and statistical details in Section 2.7; NDVI trajectories in Supplemental Figure S2). No clear phenotypic differences were visually observed among treatments. However, statistical analysis of vegetation-index-derived features identified both individual-level and group-level variation. The initial 384 vegetation-index features were reduced to 240 features by removing the two per-index extrema (minimum and maximum), the per-index undefined-pixel fraction, and any remaining zero-variance features; no feature exceeded the 50% missing-value threshold.

At the individual plant level, Plant47 (Group 7) was the most divergent sample from the control under the control-mean *t*-test, with 106 of 240 features significant at raw *p <* 0.05 and 22 significant after Benjamini–Hochberg correction (mean absolute effect size = 3.64). Under the robust median/MAD-based test, Plant32 (Group 2) was the most divergent individual, while Plant47 remained among the most extreme samples. Visual inspection of eight representative retained frames confirmed that Plant47 was consistently identified as a single, non-edge plant with no visible merging of neighboring plants and a progressively sparser, low-NDVI canopy (Supplemental Figure S3), supporting interpretation of Plant47 as an extreme individual response rather than an obvious segmentation artifact.

At the group level, a linear mixed-effects model compared each treatment group with the non-treated control across the 240 vegetation-index features, with treatment group and imaging day as fixed effects and plant identity as a random intercept. Group 2 showed the strongest group-level divergence, with 170 features significant at raw *p <* 0.05 and 162 features significant after Benjamini–Hochberg correction. Group 7 was the second most divergent group, with 137 raw-significant features and 106 FDR-significant features. Group 4 retained 9 FDR-significant features, whereas Groups 1, 3, 5, and 6 did not retain significant features after correction. Thus, Plant47 represented the strongest individual-level response under the control-mean test, while Group 2 showed the strongest group-level divergence.

#### 3.8.2 Case Study 2: Cold Stress Response in Maize

Case Study 2 evaluated whether the pipeline could translate to an independent maize cold stress dataset collected outside the Texas A&M APPG imaging platform. The goal of this case study was to assess whether phenotypic outputs extracted from standard RGB imagery could capture variation among maize hybrids across baseline, cold treatment, and recovery conditions. Representative outputs included RGB imagery, Local Binary Pattern (LBP) texture maps, morphological size analysis, and Modified Green Red Vegetation Index (MGRVI) maps from each maize plant (Figure 11a). These outputs show that the pipeline can process RGB imagery collected from a different crop species, experimental design, stress treatment protocol, and imaging system without case-specific manual intervention beyond the initial configuration. The extracted features captured variation before cold treatment, immediately following cold treatment, and after recovery. The first two principal components explained 78.8% of the total variance, with PC1 explaining 46.7% and PC2 explaining 32.1% (Figure 11b). In the score plot, point color denotes genotype and marker shape denotes treatment stage. The plot showed partial separation by timepoint: recovery samples shifted toward positive PC1 and negative PC2 values, whereas cold treatment samples shifted toward positive PC2 and showed the greatest spread along PC1, and baseline (before treatment) samples occupied an intermediate region. Genotypes were distributed within these stage clusters rather than forming fully distinct groups, with the most divergent hybrid (G1) separating from the rest along PC1. The loading analysis showed that RGB-derived vegetation indices and their summary statistics, including ExG, ExR, GLI, MGRVI, ExGR, and BIXS, contributed to this separation, captured through both their mean levels and their within-plant dispersion (standard deviation and quartiles). These results show that the pipeline extracted biologically relevant RGB-derived image features from the maize cold stress dataset, partially separated maize hybrids, and captured variation associated with cold stress and recovery. The indices driving this separation, particularly ExG and MGRVI, reflect changes in chlorophyll content and green pigment balance that are consistent with reduced photosynthetic capacity under cold stress and subsequent recovery. The outlier behavior of G1 across treatment stages may reflect a constitutively distinct pigmentation profile or a stronger physiological response to cold relative to the other hybrids.

**Figure 11:**
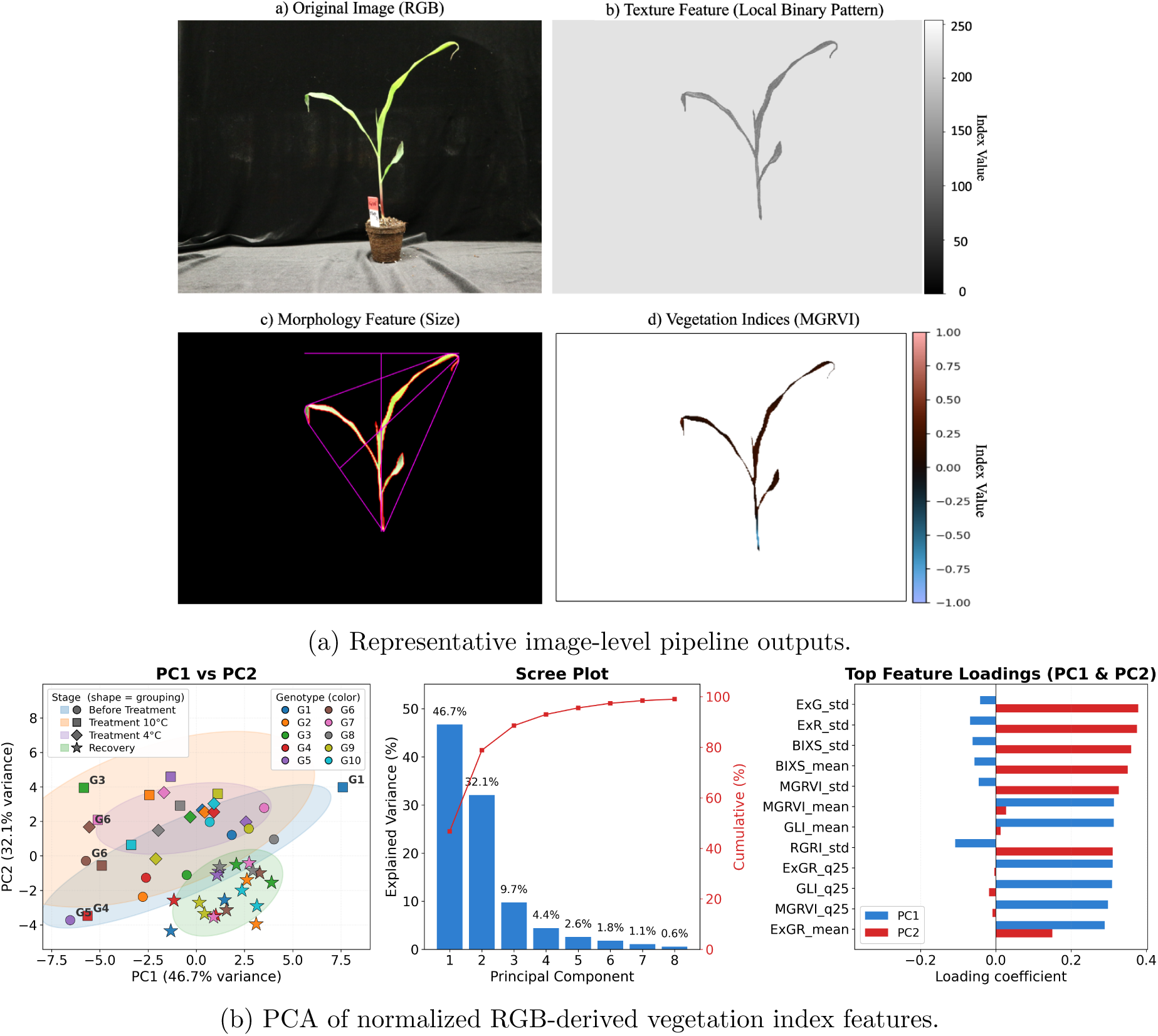
Pipeline analysis for the University of Nebraska–Lincoln CERCA maize cold stress case study. (a) Representative image-level outputs generated by the pipeline, including RGB imagery, Local Binary Pattern (LBP) texture mapping, morphological size analysis, and Modified Green Red Vegetation Index (MGRVI) visualization. (b) PCA visualization of normalized RGB-derived vegetation index features across baseline, cold treatment, and recovery samples. Point color indicates genotype, and marker shape indicates treatment stage.

## 4 Conclusion

This work provides an integrated framework for multispectral plant phenotyping within controlled environments. The primary focus and target were past and future plant science projects within the Texas A&M AgriLife Research APPG. However, the pipeline was designed with the intention that other controlled-environment facilities could adopt it, with minimal technical expertise, to extract diverse and detailed plant image features for further study, as demonstrated in the second case study.

The analytical pipeline integrates automated image acquisition, calibration, segmentation, temporal tracking, and feature extraction into a coherent and easy-to-use workflow for trait extraction under standardized environmental conditions. By combining pseudo-RGB generation with SAM v3 as the default segmentation model, BiRefNet as an alternative configurable segmentation option, and SAM2Long as the temporal instance tracking module when cross-frame identity propagation is required, the framework supports accurate plant isolation and stable identity assignment across vertically stacked image sequences. The incorporation of selfsupervised keypoint detection further enhances quantitative characterization of plant height, leaf arrangement, and canopy structure.

The Plant Growth and Phenotyping dataset version 2 (PGP v2), resulting from early APPG experiments, expands earlier efforts by providing more than 50,000 multispectral images across multiple crops and treatment groups, offering a high-quality resource for developing and benchmarking computer vision methods in controlled-environment phenomics. The two case studies demonstrated that the framework could capture both treatment-level temporal variation and stress-response phenotypes. In the maize cold stress case study, the pipeline extracted biologically relevant RGB-derived image features from an independent imaging system, partially separated genotypes, and captured variation associated with cold treatment and recovery. The organizational structure enabled standardized metadata practices and version-controlled analytical modules while maintaining biological relevance throughout development. This management philosophy builds upon the interdisciplinary model demonstrated in the Texas A&M AgriLife UAV phenotyping program, which emphasized coordinated program oversight and cross-disciplinary integration (Shi et al., 2016). The resulting datasets and workflows support the broader goal of saturating the phenome through sustained, data-driven plant science (Murray et al., 2022). Together, the dataset, analytical tools, and organizational framework provide a scalable model for future controlled-environment phenotyping initiatives. By integrating spectral, structural, and temporal information within a unified workflow, the analytical pipeline supports the broader goal of advancing transparent, data-driven, and sustainable digital agriculture. Beyond this technical pipeline, this work underscores the importance of coordinated program management and sustained collaboration among engineers, computer-vision researchers, and plant scientists. Future imaging sessions in the APPG will incorporate calibrated reference targets to enable standardized radiometric calibration across experiments and facilities, likely further improving the consistency of spectral measurements. Additionally, adjustment of the imaging head to individual plant heights is currently performed manually, representing a present limitation that will be addressed in future hardware development. Future work will explore temporal modeling for anomaly detection in longitudinal plant growth trajectories (Nia et al., 2026) and integrate neighborhood-aware mechanisms, including Neighborhood Feature Pooling (NFP) (Nia et al., 2025), to strengthen texture-aware modeling for plant stress and treatment differentiation. Future work by domain investigators could determine which extracted features are most robust for measuring cold stress and predicting stress response before visible symptoms occur. Applying this framework to additional case studies focused on quantifying genotype-dependent growth responses and treatment effects under controlled greenhouse conditions will further demonstrate the usefulness of the pipeline for advancing plant science, crop improvement, and agriculture.

## Acknowledgments

This research was supported by Texas A&M AgriLife Research. Portions of this research were conducted using the advanced computing resources provided by Texas A&M High Performance Research Computing. The authors also thank Yash Zambre, Akshatha Mohan, Omar Khater, Myla Moore, Fidel Gonzalez Torralva, and Sophia Martinez-Badillo for their valuable support and contributions to this research. The sorghum phenotyping work is supported by the Pacific Northwest National Laboratory.

## Conflict of Interest

The authors declare no conflict of interest.

## Data Availability

The Plant Growth and Phenotyping version 2 (PGP v2) dataset used in this study includes multispectral and pseudo-RGB images of maize, cotton, rice, and sorghum collected under controlled greenhouse conditions between 2023 and 2025. The dataset and associated metadata are available through the project website:

https://advanced-vision-and-learning-lab.github.io/PGP_Website/. The complete pipeline implementation, including preprocessing, segmentation, tracking, stitching, feature extraction, and downstream analysis, is publicly available through the Plant Analysis Tool Pipeline repository:

https://github.com/Advanced-Vision-and-Learning-Lab/Plant_Analysis_Tool_Pipeline.

## Abbreviations

AI: Artificial Intelligence
BEN: Background Erase Network
BiRefNet: Bilateral Reference Network
CNN: Convolutional Neural Network
EHD: Edge Histogram Descriptor;
NDVI: Green Normalized Difference Vegetation Index
HOG: Histogram of Oriented Gradients
L: Lacunarity
LBP: Local Binary Pattern
ML: Machine Learning
MSIS: Multispectral Imaging System
NIR: Near-Infrared (spectral band)
NDVI: Normalized Difference Vegetation Index
PGP: Plant Growth and Phenotyping
RE: Red-Edge (spectral band)
RGB: Red-Green-Blue composite image
ROI: Region of Interest
SAM: Segment Anything Model
SAM2Long: Segment Anything Model for Long Sequences
SDK: Software Development Kit
YOLO: You Only Look Once (object detection model)
UAV: Unmanned Aerial Vehicle
BRISK: Binary Robust Invariant Scalable Keypoints
ORB: Oriented FAST and Rotated BRIEF
SIFT: Scale-Invariant Feature Transform
AKAZE: Accelerated KAZE;

## Notes

### Competing Interest Statement

The authors have declared no competing interest.

### Summary of Updates

This version of the manuscript has been revised to incorporate a second case study and include additional data. Furthermore, the supplementary materials have been separated into distinct files to improve clarity and organization.

