## Supplemental Material for "A Data-Driven Image Extraction and Analysis Pipeline for Plant Phenotyping in Controlled Environments"

### Supplemental File

This supplemental file contains six pages, three supplemental figures, and three supplemental tables.

This supplemental material provides additional figures, tables, and supporting information for the manuscript titled *A Data-Driven Image Extraction and Analysis Pipeline for Plant Phenotyping in Controlled Environments.*

### Metadata

The greenhouse imaging system records structured metadata for each image, including plant identity, spatial location, acquisition time, and robotic positioning parameters. Spatial coordinates are reported in millimeters (mm), and timestamps are reported in local system time. A categorical orientation variable indicates the predefined camera viewpoint. This metadata enables reliable plant identification, spatial calibration, and temporal tracking across imaging sessions.

Supplemental Table S1: Example metadata recorded during image acquisition. Img denotes image index; Ori. denotes camera orientation, representing predefined viewpoints; PX and PY represent plant spatial coordinates in mm; RX, RY, and RZ represent robot spatial coordinates in mm.

| Plant ID | Room | Date | Time | Img | Row | Ori. | PX | PY | RX | RY | RZ |
|---|---|---|---|---|---|---|---|---|---|---|---|
| Cotton 1 | 2 | 2023-05-16 | 13:01 | 1 | 1 | 2 | 1930 | 950 | 1930.0 | 600 | 925.0 |
| Cotton 1 | 2 | 2023-05-16 | 13:01 | 2 | 1 | 2 | 1930 | 950 | 1930.0 | 600 | 980.0 |
| Cotton 2 | 2 | 2023-05-16 | 13:01 | 1 | 1 | 2 | 2630 | 950 | 2630.0 | 600 | 925.0 |
| ... | | | | | | | | | | | |

### Qualitative Segmentation Comparison

This figure provides a qualitative comparison of classical segmentation methods, including Otsu thresholding and watershed segmentation, versus SAM v3 across four crop species. This comparison complements the quantitative segmentation results reported in the main manuscript.

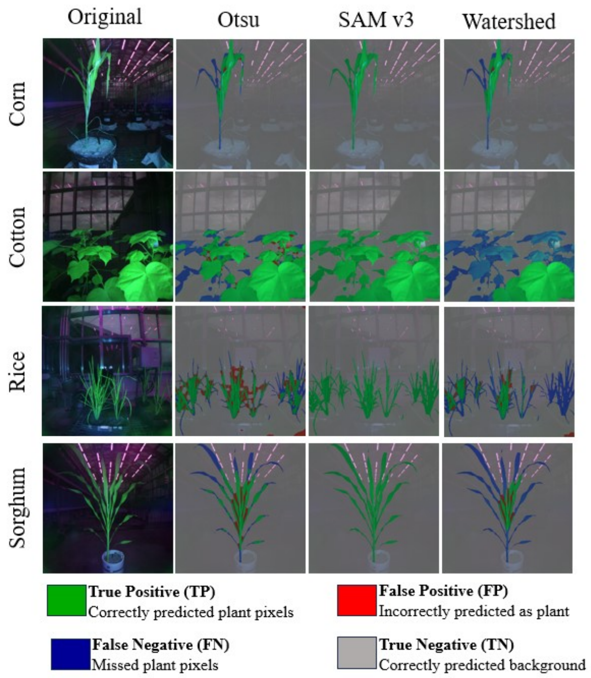


Supplemental Figure S1: Qualitative comparison of classical segmentation methods versus SAM v3 across four crop species. SAM v3 produced more accurate segmentation masks across species.

### Vegetation Indices

The following vegetation indices were computed from the multispectral bands of the images to characterize plant health, canopy structure, and reflectance properties. These indices include common vegetation metrics, such as the Normalized Difference Vegetation Index (NDVI), Green NDVI (GNDVI), Red-Edge NDVI (NDRE), and others that capture different aspects of plant physiology. The formulas for each index are provided in Supplemental Table S2, which is split into two parts for clarity.

Supplemental Table S2: Complete list of vegetation indices computed in the pipeline.

| # | Index | Full name | Formula |
|---|---|---|---|
| 1 | NDVI | Normalized Difference Vegetation Index | $\frac{NIR-Red}{NIR+Red}$ |
| 2 | GNDVI | Green NDVI | $\frac{NIR-Green}{NIR+Green}$ |
| 3 | NDRE | Red-Edge NDVI | $\frac{NIR-RE}{NIR+RE}$ |
| 4 | GRNDVI | Green–Red NDVI | $\frac{NIR-(Green+Red)}{NIR+(Green+Red)}$ |
| 5 | TNDVI | Transformed NDVI | $\sqrt{\frac{NIR-Red}{NIR+Red}}+0.5$ |
| 6 | MGRVI | Modified Green–Red VI | $\frac{Green^2-Red^2}{Green^2+Red^2}$ |
| 7 | GRVI | Green Ratio VI | $\frac{NIR}{Green}$ |
| 8 | NGRDI | Normalized Green–Red Difference | $\frac{Green-Red}{Green+Red}$ |
| 9 | MSAVI | Modified SAVI | $0.5\Big(2NIR+1 - \sqrt{(2NIR+1)^2-8(NIR-Red)}\Big)$ |
| 10 | OSAVI | Optimized SAVI | $\frac{NIR-Red}{NIR+Red+0.16}$ |
| 11 | TSAVI | Transformed SAVI | $\frac{s(NIR-sR-a)}{aNIR+R-as+X(1+s^2)}$ |
| 12 | GSAVI | Green SAVI | $1.5\frac{NIR-Green}{NIR+Green+0.5}$ |
| 13 | GOSAVI | Green Optimized SAVI | $\frac{NIR-Green}{NIR+Green+0.16}$ |
| 14 | GDVI | Green Difference VI | $NIR-Green$ |
| 15 | NDWI | Water Index | $\frac{Green-NIR}{Green+NIR}$ |
| 16 | CIRE | Chlorophyll Index Red-Edge | $\frac{NIR}{RE}-1$ |
| 17 | LCI | Leaf Chlorophyll Index | $\frac{NIR-RE}{NIR+RE}$ |
| 18 | $CI_{green}$ | Chlorophyll Index Green | $\frac{NIR}{Green}-1$ |
| 19 | MCARI | Modified Chlorophyll Absorption Ratio Index | $\big((RE-Red)-0.2(RE-Green)\big)\cdot\frac{RE}{Red}$ |
| 20 | MCARI1 | MCARI Variant 1 | $1.2[2.5(NIR-Red)-1.3(NIR-Green)]$ |
| 21 | MCARI2 | MCARI Variant 2 | $\frac{1.5[2.5(NIR-Red)-1.3(NIR-Green)]}{\frac{1}{\sqrt{(2NIR+1)^2-(6NIR-5\sqrt{Red})}}}$ |
| 22 | MTVI1 | Modified TVI 1 | $1.2[1.2(NIR-Green)-2.5(Red-Green)]$ |
| 23 | MTVI2 | Modified TVI 2 | $\frac{1.5[1.2(NIR-Green)-2.5(Red-Green)]}{\frac{1}{\sqrt{(2NIR+1)^2-(6NIR-5\sqrt{Red})-0.5}}}$ |
| 24 | CVI | Chlorophyll Vegetation Index | $\frac{NIR\cdot Red}{Green^2}$ |
| 25 | ARI | Anthocyanin Reflectance Index | $\frac{1}{Green}-\frac{1}{RE}$ |
| 26 | ARI2 | Anthocyanin Reflectance Index 2 | $NIR\left(\frac{1}{Green}-\frac{1}{RE}\right)$ |
| 27 | DVI | Difference Vegetation Index | $NIR-Red$ |
| 28 | WDVI | Weighted DVI | $NIR-0.5Red$ |
| 29 | SR | Simple Ratio | $\frac{NIR}{Red}$ |
| 30 | MSR | Modified Simple Ratio | $\frac{(NIR/Red)-1}{\sqrt{(NIR/Red)+1}}$ |
| 31 | PVI | Perpendicular VI | $\frac{NIR-0.5Red-0.3}{\sqrt{1+0.5^2}}$ |
| 32 | EVI2 | Two-band EVI | $\frac{2.4(NIR-Red)}{NIR+Red+1}$ |
| 33 | DSWI4 | Disease Stress Water Index 4 | $\frac{Green}{Red}$ |
| 34 | GEMI | Global Environmental Monitoring Index | $\eta(1-0.25\eta)-\frac{Red-0.125}{1-Red}$, $\eta=\frac{2(NIR^2-Red^2)+1.5NIR+0.5Red}{NIR+Red+0.5}$ |
| 35 | ExR | Excess Red Index | $1.3\,Red-Green$ |
| 36 | RI | Redness Index | $\frac{Red-Green}{Red+Green}$ |
| 37 | RRI | Red-Edge Ratio Index | $\frac{NIR}{RE}$ |
| 38 | RRI1 | Red-Edge Ratio Index 1 | $\frac{NIR}{RE}$ |
| 39 | RRI2 | Red-Edge Ratio Index 2 | $\frac{RE}{Red}$ |
| 40 | AVI | Advanced Vegetation Index | $\sqrt[3]{NIR(1-Red)(NIR-Red)}$ |
| 41 | SIPI2 | Structure-Insensitive Pigment Index 2 | $\frac{NIR-Green}{NIR-Red}$ |
| 42 | TCARI | Transformed Chlorophyll Absorption Ratio Index | $3\Big[(RE-Red) - 0.2(RE-Green)\frac{RE}{Red}\Big]$ |
| 43 | TCARIOSAVI | TCARI/OSAVI | $\frac{3(RE-Red)-0.2(RE-Green)\frac{RE}{Red}}{\frac{1}{1+0.16\frac{NIR-Red}{NIR+Red+0.16}}}$ |
| 44 | CCCI | Canopy Chlorophyll Content Index | $\frac{(NIR-RE)(NIR+Red)}{(NIR+RE)(NIR-Red)}$ |
| 45 | RDVI | Renormalized Difference Vegetation Index | $\frac{NIR-Red}{\sqrt{NIR+Red}}$ |
| 46 | NLI | Nonlinear Vegetation Index | $\frac{NIR^2-Red}{NIR^2+Red}$ |
| 47 | BIXS | Brightness Index (Squared) | $\sqrt{\frac{Green^2+Red^2}{2}}$ |
| 48 | IPVI | Infrared Percentage Vegetation Index | $\frac{NIR}{NIR+Red}$ |

### Inventory of Extracted Features

This section summarizes the complete feature inventory generated by the APPG multispectral phenotyping pipeline. The 863-dimensional feature vector integrates spectral vegetation indices, multi-domain texture descriptors, and detailed morphological traits, providing a comprehensive quantitative representation of plant reflectance, structure, and spatial heterogeneity for downstream statistical and machine learning analyses.

Supplemental Table S3: Breakdown of the 863-dimensional feature vector generated by the APPG multispectral phenotyping pipeline.

| **Feature family** | **Base names / components** | **Count** |
|---|---|---|
| Vegetation indices | **Stats (8):** mean, std, min, max, median, q25, q75, nan_fraction.<br>**Indices (48):** NDVI, GNDVI, NDRE, GRNDVI, TNDVI, MGRVI, GRVI, NGRDI, MSAVI, OSAVI, TSAVI, GSAVI, GOSAVI, GDVI, NDWI, DSWI4, CIRE, LCI, CIgreen, MCARI, MCARI1, MCARI2, MTVI1, MTVI2, CVI, ARI, ARI2, DVI, WDVI, SR, MSR, PVI, GEMI, ExR, RI, RRI1, RRI2, RRI, AVI, SIPI2, TCARI, TCARIOSAVI, CCCI, RDVI, NLI, BIXS, IPVI, EVI2. | 384 |
| Texture descriptors | **Bands (6):** color, nir, red_edge, red, green, pca.<br>**Blocks (6):** lbp, hog, lac1, lac2, lac3, ehd_map.<br>**Stats (5):** mean, std, min, max, median. | 180 |
| Texture (EHD channels) | **Bands (6):** color, nir, red_edge, red, green, pca.<br>**Channels (9):** channel_0 to channel_8.<br>**Stats (5):** mean, std, min, max, median. | 270 |
| Morphology (scalar traits) | **Scalar traits (25):** area, area_cm2, aspect_ratio, bbox_area_cm2, circularity, convex_hull_area, convex_hull_vertices, convexity, ellipse_angle, ellipse_eccentricity, ellipse_major_axis, ellipse_minor_axis, elongation, height, height_cm, longest_path, major_axis_cm, minor_axis_cm, num_leaves, num_stems, perimeter, perimeter_cm, solidity, width, width_cm. | 25 |
| Morphology (tuple components) | **Tuples:** center_of_mass, ellipse_center.<br>**Axes:** x, y. | 4 |
| **Total** | | **863** |

### Case Study 1

The sorghum case study included 48 plants assigned to seven LEEB mutagenesis treatment groups (G1–G7) and one non-treated control group (NT). NDVI trajectories were automatically extracted across imaging dates. While overall trends were similar, differences in magnitude and temporal dynamics were observed across treatments. For comparison, NDVI values were normalized to the control baseline at each date. The resulting difference curves highlight divergence from the control over time, demonstrating the pipeline's ability to capture structured variation among treatments.

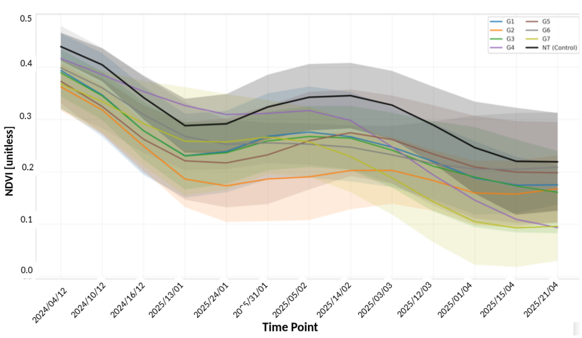


(a) Mean NDVI trajectories, reported as mean ± SD, for all treatments across fourteen imaging dates.

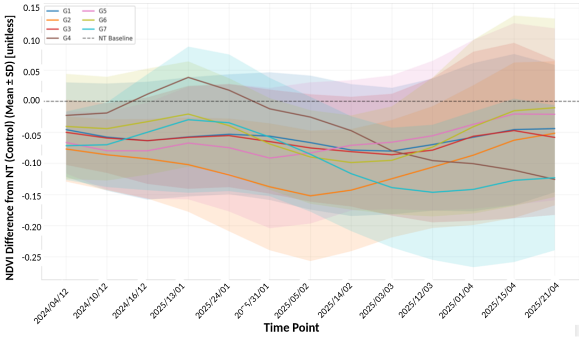


(b) NDVI differences relative to the NT baseline, reported as mean ± SD.

Supplemental Figure S2: Temporal NDVI analysis for the mutagenized sorghum case study.

Prior to statistical testing, every plant-date segmentation was screened, and frames with occlusion, mis-segmentation, or non-detection were excluded. For the most divergent individual, Plant47, inspection of eight representative retained frames (Figure S3) confirmed a single, consistently identified, non-edge plant with no merging of neighboring plants and a steadily declining NDVI canopy, indicating a genuine individual response rather than a segmentation artifact.

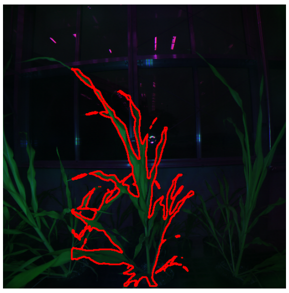

(a) 2024-12-04

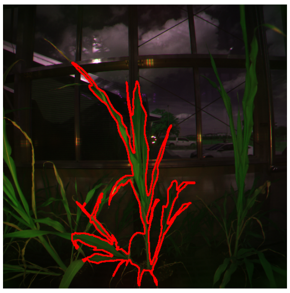

(b) 2024-12-16

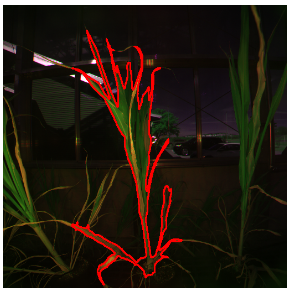

(c) 2025-01-13

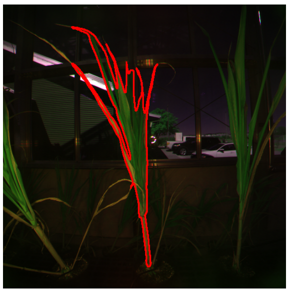

(d) 2025-01-31

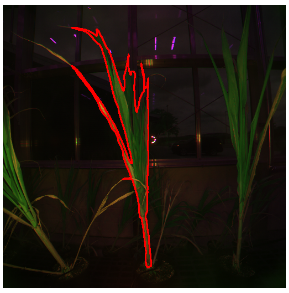

(e) 2025-02-05

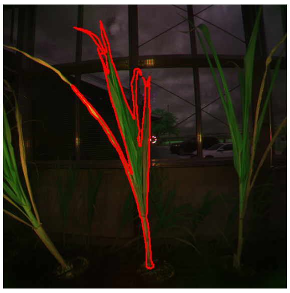

(f) 2025-03-03

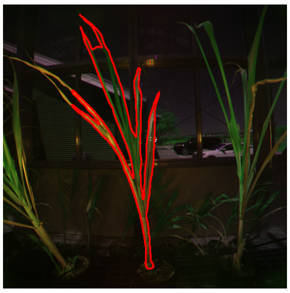

(g) 2025-03-24

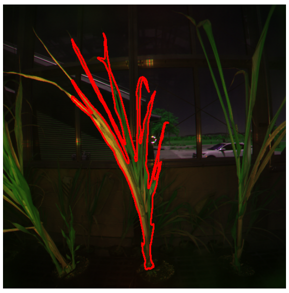

(h) 2025-04-21

Supplemental Figure S3: Visual verification of Plant47 segmentation quality using eight representative retained frames. Panels show Plant47 with the segmentation mask outlined in red, confirming that the plant was consistently identified as a single, non-edge individual with no merging of neighboring plants and a declining canopy pattern.
